# Brain precapillary sphincters modulate myogenic tone in adult and aged mice

**DOI:** 10.1101/2025.04.11.648315

**Authors:** Christina Fjorbak, Nikolay P. Kutuzov, Teddy Groves, Martin Lauritzen, Søren Grubb

## Abstract

Brain precapillary sphincters, which are surrounded by contractile pericytes and are located at the junction of penetrating arterioles and first-order capillaries, can increase their diameter by ∼30% in a few seconds during sensory stimulation, allowing for rapid control of capillary blood flow over a wide dynamic range. We hypothesized that these properties could help precapillary sphincters maintain the capillary blood flow and shield the downstream capillaries during surges in blood pressure. To test this, we visualized microvessels in adult and old anaesthetized mice using *in vivo* two-photon microscopy. We showed that a blood pressure surge disrupts both microvascular myogenic response and neurovascular coupling in both adult and old mice, with old mice exhibiting a more diminished myogenic response. Similarly, laser ablation of contractile pericytes encircling precapillary sphincters disrupted neurovascular coupling and myogenic response. The resistance provided by precapillary sphincters may be increasingly important in old mice, where we found changes in the topology of microvessels, potentially affecting microvascular blood flow. Old mice displayed more tortuous penetrating arterioles, reduced pial collateral arteriolar density and altered capillary densities: reduced in the arterial end and increased in the venous end. Our results illustrate how blood pressure surges affect brain microvascular function, underscore the protective role of precapillary sphincters during cerebrovascular autoregulation in response to blood pressure surges and compare vascular topology in adult and old mice *in vivo*.

## Introduction

A brain precapillary sphincter (PS) – a narrowing of the vasculature surrounded by layers of contractile pericytes – connects a penetrating arteriole (PA) to a first-order capillary [1–7]. This placement of PSs makes them key players in the regulation of cerebral blood flow [1–3,8]. PSs can both increase and decrease capillary blood flow over a wide dynamic range, made possible by large relative diameter changes of PSs: ∼30-60% dilation and ∼80% constriction [1,7]. Therefore, we hypothesized that PSs could help maintain blood flow and shield downstream capillaries during brief and rapid increases in systemic blood pressure (BP) [9,10].

The primary mechanism of autoregulation is the myogenic Bayliss response: a vascular contraction or dilation in response to high or low blood pressure, respectively [11]. The myogenic response, triggered by variations in intraluminal pressure, is compromised by aging and high BP fluctuations [10,12]. Here, we examined the PS responses to abrupt, high amplitude increases in BP to determine whether PSs help to preserve the myogenic response, as recently suggested in a computational study [8].

Preserving the myogenic response in ageing is crucial because age-related arterial stiffening, combined with hypertension, increase the amplitude of the pulse pressure (PP) waves, allowing them to penetrate deep into the cerebral microcirculation [9,10,13–15]. This exposes microvessels to mechanical stress, causing disruption of the neurovascular unit, microbleeds, loss of microvessels and cognitive decline [14,16–19]. The thin capillary walls are susceptible to mechanical damage because of less coverage by pericytes, compared to larger microvessels, and lack of elastin and collagen fibrils [4]. We hypothesized that the PS, placed at the entrance to the capillary bed, can dampen PP; thus, protecting the downstream vasculature from pulsatile stress. To investigate this, we developed a new method for analysis of PP-driven pulsations of microvessels diameters.

With aging, the brain vasculature becomes less responsive to vasoactive molecules, impairing its ability to maintain steady basal blood flow and to rapidly change the blood flow [3,20,21]. This is accompanied by a decrease in vessel density near PAs and by a loss of pericyte processes, although the number of pericyte somas and their α-smooth-muscle actin density are preserved [3]. In succession to our previous study of microvascular topology with *in vivo* two-photon microscopy [3], here we studied four times larger brain volumes (∼0.5 mm^3^ per mouse), containing several PAs and ascending venules. We hypothesized that aging-related changes in capillary density depend on the capillary order, and that the changes in microvascular topology might affect PS regulation of the BP and PP in the capillary bed, especially during acute BP increases, reducing its ability to protect the downstream capillaries.

## Materials and methods

### Animal handling

The Danish National Ethics Committee approved all animal procedures, which followed the European Council’s Convention for the Protection of Vertebrate Animals Used for Experimental and Other Scientific Purposes. We adhered to all applicable ethical standards for animal research. A total of 74 male or female NG2-dsRed (Tg(Cspg4-DsRed.T1)1Akik/J; Jackson Laboratory) or C57bl6/j were used: 46 adults (15-30 weeks old) and 28 old (90-110 weeks old).

### Surgical procedures

Anesthesia was initiated with xylazine (10 mg/kg i.p.) and ketamine (60 mg/kg i.p.), with additional ketamine doses (30 mg/kg i.p.) during surgery. Lidocaine (0.15 mL, 5 mg/mL) was given subcutaneously for local pain relief. We performed tracheotomy to mechanically ventilate mice (Minivent type 845, Harvard Apparatus; Fig. 1a). Three catheters were inserted: first catheter – in the left femoral artery to monitor blood pressure, second catheter – in the left femoral vein to continuously supply a mixture of FITC-dextran and α-chloralose (25% w/vol; 0.02 mL/10 g/h), and third catheter – in the right femoral vein to inject angiotensin II (0.25µg/kg/min) (Fig. 1a). For some experiments, we took arterial blood samples (50 µL) to measure partial pressures of O_2_ (pO_2_) and CO_2_ (pCO_2_), as well as pH in the blood using an ABL700 (Radiometer, Copenhagen). The measured pO_2_ and pCO_2_ were within 95–110 mmHg and 35–40 mmHg, respectively; pH was between 7.35–7.45. Adjustments to the respiration rate and the composition of the air supply were made based on the recorded pO_2_ and pCO_2_ and continuous end-expiratory CO_2_ monitoring using a Capnograph 340 (Harvard Apparatus). A head bar was affixed and a craniotomy (diameter of ∼3 mm, located 3 mm to the right and 0.5 mm behind the bregma) was performed over the right somatosensory barrel cortex. An acute cranial window was established using an agarose solution made with artificial cerebrospinal fluid and a glass coverslip glued to the skull. Body temperature was maintained at 37°C with a feedback-controlled heating pad. After surgery, the anesthesia was switched to α-chloralose (25% w/vol; 0.02 mL/10 g/h). Post-experiment, mice were euthanized with a pentobarbital injection followed by cervical dislocation.

**Fig. 1.**
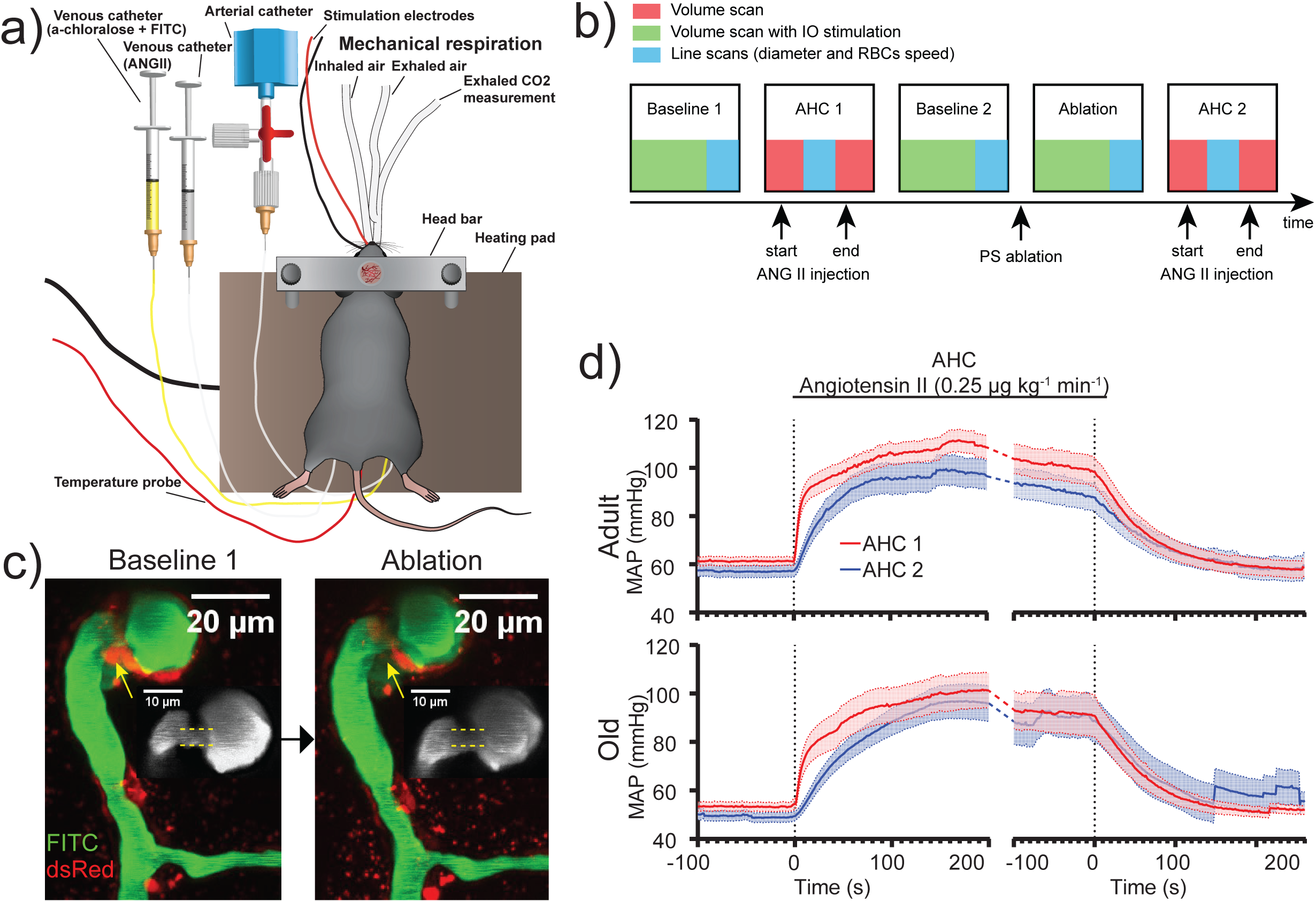
Experimental pipeline for in vivo two-photon microscopy and acute hypertensive challenge: **a**) Experimental setup for *in vivo* two-photon microscopy. **b**) Our experiment consisted of five groups of measurements, including 3D volume scans and 1D line scans, during which we applied five *treatments* (boxes). “Baseline 1” is the first control (i.e. no treatment); “AHC 1” – blood pressure increase after injection of angiotensin II (ANG II); “Baseline 2” – second control, after the blood pressure returns back to the *Baseline 1* level; “Ablation” – laser ablation of the PS; “AHC 2” – blood pressure increase after the second injection of ANG II. **c**) Maximum intensity projection of a volume stack of images, taken at different depths, of pericytes (NG2-dsRed, red) and blood vessels (FITC-dextran, green) around a PS before (left) and after (right) laser ablation of the pericytes encircling the PS (yellow arrow). **Insets** (gray): a zoomed-in view of FITC-dextran fluorescence around the PS, showing its dilation after the ablation. **d**) Mean arterial pressure (MAP) during AHC 1 (red) and AHC 2 (blue) in adult and old mice. For comparison, we aligned MAP traces from both AHC measurements such that the rise of MAP begins at ∼0 sec (left part of each graph); Similarly, we aligned the traces of MAP decrease (after the stop of ANG II injection), such that the MAP starts to decrease at ∼0 sec (right part of each graph). Some old mice died ≥ 130 s after the ANG II infusion stop during AHC 2 and their MAP traces, showing zero MAP after the death, were excluded from the calculation of the mean traces. This shows as an artefact – a jump in MAP ∼130 s after ANG II infusion stops.

### Whisker pad stimulation (IO stimulation)

We used electrical whisker pad stimulation to activate the mouse’s sensory barrel cortex. Custom-made bipolar electrodes, inserted through the skin, stimulated the contralateral ramus infraorbitalis (IO) of the trigeminal nerve. The cathode was positioned near the hiatus IO and the anode inserted into the masticatory muscles (Fig. 1a). Thalamocortical IO stimulation was carried out with an intensity of 1.5 mA (using ISO-flex by A.M.P.I.) for 1 ms, delivered in 20-second trains at a frequency of 2 Hz.

### Acute hypertensive challenge

Acute hypertensive challenge (AHC) was induced by injecting 4.8 µM angiotensin II, diluted in saline, via the right femoral vein catheter (0.25 µg/kg/min, Fig. 1a). AHC 1 and AHC 2 lasted ∼8 min.

### Two-photon imaging: Setup

We used a two-photon microscope (FluoView FVMPE-RS, Olympus) with a 25x (1.05 NA) water-immersion objective (Olympus). To maximize the quality of recorded images and facilitate data analysis, we removed dark output and periodic bias from the detectors (PMTs) using the “quick-and-noisy” method [22]. We illuminated the brain with 920 nm laser light and collected fluorescence in (1) the 510-560 nm range (“green channel”) from FITC-dextran labeling blood vessels and in (2) the 590-650 nm range (“red channel”) from dsRed labeling pericytes and smooth muscle cells. We observed autofluorescent debris in the parenchyma of old mice (Fig. 6c, NG2-dsRed), which were likely due to lipofuscin deposits [23]. This autofluorescence appeared in both green and red channels (though more in red), which we used to discriminate it from the NG2-dsRed fluorescence, which appeared only in the red channel.

### Two-photon imaging: Recording volumes in time (xyzt)

We selected the volume for imaging by choosing a PS on a PA, from which one can easily trace branching capillaries down to the fifth order. This PA and five orders of capillaries originating from it we denote as a vessel tree. We recorded one vessel tree per mouse. PSs were detected as described below in “Quantification of precapillary sphincters and bulbs”. We selected only arterioles whose cylinder-axes were roughly parallel with the fixed scanning direction of the resonant scanner. Images (512x512 pixels, 1 µm/pixel) were taken at different depths, with 2 µm steps, to track the PA from the pial artery to a depth of 500-600 µm. Higher-resolution images (512x512 pixels, 0.2 µm/pixel, with 1 µm steps in depth) were taken over a depth of 20-40 µm, centered at the PS and the initial 2-3 orders of capillaries. Volumetric imaging in time (xyzt) of PSs and up to the 2^nd^ order of capillaries were performed during IO stimulation and AHCs (Fig. 1b).

### Two-photon imaging: Recording kymograms – line-scans in time (xt)

Kymograms, i.e. line-scans in time, used to estimate red blood cell (RBC) velocity, RBC_v_, in the PSs and 1^st^-order capillaries, were recorded by bi-directional resonance scanning along the vessel’s cylinder-axis at 512 pixels per line, 0.124 µm/pixel, and 16 kHz sampling rate. Kymograms, used to estimate capillary diameter pulsations or RBC velocity downstream of the 1^st^-order capillary, were recorded by uni-directional galvo-scanning along a line perpendicular to the vessel’s cylinder-axis, i.e. across the vessel at 25-100 pixels per line, 0.199 µm/pixel, and 0.8 kHz sampling rate.

### Precapillary sphincter ablation

To photoablate PSs [24], we increased the laser intensity 4-8 times while line-scanning along the somas of the pericytes encircling the PSs (Fig. 1c), guided by the NG2-DsRed fluorescence and unguided for WT mice. We considered the ablation completed when the PS dilated following the ablation as judged by eye (Fig. 1c). If the PS did not dilate, we continued ablating until it did.

Occasionally, the ablation caused a leakage of FITC into the space surrounding the PS and 1^st^-order capillary (Supplementary Fig. 1 and Video 3). These data were discarded.

### Blood pressure data

Arterial blood pressure was measured via a femoral artery catheter and sampled at 0.2 kHz in SPIKE2.7 via a BP-1 pressure monitor (World Precision Instruments). Mean arterial pressure (MAP) was calculated as the average blood pressure over the cardiac cycle using SPIKE2.7. Note that MAP following surgery and shift to α-chloralose was bordering hypotension (<60 mmHg) in both adult and old mice (Adult: 63.5±2.1 mmHg, Old: 53.4±1.9 mmHg, Table. 1). This is not uncommon for anaesthetized mice, as MAPs have been reported within a range of 51±3 to 95±9 mmHg [25]. During AHC, the MAP increased within the initial 10-20 seconds (Fig. 1d), in both adult and old mice (Table 1).

**Table 1:** Mean arterial blood pressure (MAP), pulse pressure (PP), and the heart rate (HR) in adult and old mice presented as mean ± S.E.M. For AHC 1 & 2, the measurements are from the end of the angiotensin II injection.

|  | MAP (mmHg) |  | PP (mmHg) |  | HR (Hz) |  |
| --- | --- | --- | --- | --- | --- | --- |
|  | Adult | Old | Adult | Old | Adult | Old |
| Baseline 1 | 63.8 ± 2.3 | 53.4 ± 1.9 | 12.6 ± 1.5 | 9.9 ± 0.9 | 5.2 ± 0.2 | 6.1 ± 0.3 |
| AHC 1 | 115 ± 8 | 88.2 ± 8.6 | 18.3 ± 4.4 | 15.9 ± 3.5 | 5.3 ± 0.3 | 6.8 ± 0.6 |
| Baseline 2 | 60.7 ± 2.3 | 50.2 ± 2.5 | 10.4 ± 1.0 | 9.9 ± 1.1 | 5.4 ± 0.1 | 6.4 ± 0.3 |
| Ablation | 58.3 ± 2.5 | 47.9 ± 1.8 | 10.4 ± 1.0 | 9.8 ± 1.0 | 5.5 ± 0.1 | 6.2 ± 0.2 |
| AHC 2 | 92.2 ± 6.8 | 90.3 ± 8.9 | 13.7 ± 2.9 | 12.3 ± 3.3 | 5.6 ± 0.3 | 6.4 ± 0.7 |

### Vessel diameters estimated from volumetric data (xyzt)

First, the volumetric data (xyzt), a stack of images recorded at different depths (xyz) in time (t), were corrected for drift (translation in 3D) using the “3D drift correction” plugin for FIJI. Second, at each time point, the stacks of images were projected from 3D to 2D along the microscope’s objective z-axis, i.e. in depth, using maximum intensity projection (MIP). Third, from the projected images in time, we estimated diameters of the blood vessels with a custom-made MATLAB script, employing a Chan-Vese segmentation algorithm [1,3].

### Vessel diameters estimated from kymograms (xt)

First, we averaged recorded kymograms over non-overlapping blocks of 16 line-scans (∼20 ms). Second, we subtracted the average background fluorescence, estimated by averaging pixels away from the vessel, from every line-profile in the block-averaged kymogram. Third, we selected a threshold (inset in Fig. 5a), ∼10-20% of the maximal intensity of the line-profile, and calculated positions of the vessel’s left, x_1_, and right, x_2_, edges (inset in Fig. 5a) along the scan line, where the threshold intersected the line-profile. Keeping the threshold as low as possible ensured that estimated diameters of the vessels were not affected by the fluctuating fluorescence intensity in the central part of the vessel due to flowing RBCs. Fourth, we calculated the diameter, d = x_2_ -x_1_, and the center position, x_c_ = (x_1_ + x_2_)/2 (Fig.5b). We excluded data from the analysis if FITC-dextran leaked in the extravascular space (Supplementary Fig.1, Video 3) or if the blood flow stopped during the recording.

### Spectral power of pulsation of the vessel diameters, Pd, and their center positions, Pc

We calculated power spectra of d(t) and x_c_(t) using the fast Fourier transform (FFT). We minimized spectral leakage, as described in section IIIB in [22], which resulted in narrow spectral peaks (Fig. 5c). A typical power spectrum (Fig. 5c) contained (i) low-frequency components (< 2 Hz), corresponding to slow spontaneous motion of a blood vessel; (ii) a peak corresponding to the ventilation frequency (around 3.5 Hz); and (iii) a peak corresponding to the heartbeat frequency. We did not use the higher harmonics of the heartbeat pulsations. From each power spectrum we extracted spectral power of diameter, P_d_, and center, P_c_, pulsations at the heartbeat frequency. We estimated white noise floor amplitude by averaging spectral power between 13 and 15 Hz, away from any peaks, and found that the noise’s amplitude was typically 1-2 orders of magnitude lower than P_d_ or P_c_ (Fig. 5c). Therefore, we did not subtract the white-noise background from the estimated P_d_ or P_c_.

### Red blood cell velocity and flux

RBC velocity, RBC_v_, was calculated from kymograms (xt) with a Matlab script that uses hybrid image filtering and iterative Radon transformation [26]. RBC flux, RBC_flux_, was calculated by drawing a line along the kymogram using ImageJ and counting the number of intensity troughs, i.e. RBC shadows, per time using the software Spike2.

### Myogenic response analysis

We quantified the myogenic response of a vessel to AHC by estimating the Pearson’s correlation coefficient (CC) between the time series of mean arterial pressure (MAP) and the vessel’s diameter. The start of the time traces was the start of the MAP increase, which we could select by hand thanks to large increases of MAP (∼50-100%, Table 1) during AHC. The duration of the traces, from which we estimated CC, was no longer than 3 minutes, depending on the length of the vessel’s diameter trace. Some diameter traces were shorter than others, for example because the vessel’s cylinder axis moved out of the focal plane during collection of the line-scans, preventing diameter estimation.

### Quantification of precapillary sphincters and bulbs

PSs and bulbs were detected by comparing the diameter of a PS at the junction between a PA and a 1^st^ order capillary with the diameter of the 1^st^-order capillary. The branchpoint was categorized as having a PS if its diameter was <80% of the 1^st^-order capillary’s diameter and having a bulb if the diameter immediately downstream of the PS was >125% of the 1^st^-order capillary’s diameter.

### Wide-field angiograms to study topology of collaterals

We recorded wide-field fluorescence from whole craniotomies with a 4x (0.16 NA) air objective (Olympus), a CAM-ORCA-FLASH-2.8 - Scientific CMOS 2.8 Megapixel Camera (Hamamatsu), and a 100W Mercury lamp as a light source. We recorded videos of the brain surface vessels (10 Hz sampling rate) while injecting FITC-dextran in the blood and observed how it appeared first in the arterioles and then in the veins a few seconds later. From these videos, we created a map of the craniotomy (ImageJ’s time-lapse color-coding plugin), where each pixel shows the time of the arrival of FITC-dextran-labeled blood plasma (Fig. 6a, Video 4). These maps facilitated identification of pial arteries and veins (red and blue in Fig. 6a) and allowed us to examine age-related changes in pial vessels. We also used these data to analyze collaterals on the brain’s surface.

### Two-photon angiograms to study topology of capillaries

To study the topology of cortical capillaries, we recorded stacks of images at different depths (xyz) with two-photon microscopy from four adult and four old anesthetized NG2-dsRed or WT mice, with blood vessels labeled with FITC-dextran. First, we acquired a total volume of >0.5 µm^3^ consisting of nine or more ∼25% overlapping >500 µm deep volume stacks throughout the entire cranial window, ensuring that the volumes analyzed were comparable between adult and old mice (Video 5). Second, individual stacks were stitched together in FIJI using the “Stitching” plugin. Third, we analyzed the stitched stacks using Amira software (Thermo Fisher), e.g., using despeckle- and Gaussian filters, Z-drop corrections, and thresholding. Fourth, we skeletonized the vascular lumen (green channel) and manually labelled large vessels and capillaries. The labeling was limited to 12 orders of capillaries (1^st^ order capillaries branch from PAs), as this accounted for most of the capillary bed.

### Statistics: Bayesian multilevel models

The data analyzed have a nested structure, i.e. several measurements are taken from a given vessel in each mouse. As a result, when comparing data pooled from different mice, data recorded from individual mice can be correlated (not independent). This prevented us from using standard statistical tests, e.g., t-test or ANOVA, that assume independence of the measurements.

Data with nested structure can be analyzed with a frequentist [27] or Bayesian [28,29] approaches. We chose Bayesian multilevel generalized linear regression, which allows to incorporate non-experimental information (e.g. data collection features and data properties) into the statistical analysis for more efficient analysis of small data sets (<10 measurements). Multilevel generalized linear models (GLMs) provide a powerful framework for modelling structured data [30,31].

Bayesian multilevel GLMs allow encoding information about latent parameters using prior distributions, thereby conferring several advantages over non-Bayesian approaches including regularization, computational tractability and model identification among others [32]. Bayesian multilevel GLMs have successfully been applied to many similar problems [33,34].

All our Bayesian multilevel GLMs are structurally similar but differ in their measurement distributions, parameter dependencies, and prior distributions. These differences arose organically: in each case we started with a simple, naive model of the target measurement type, then iteratively added and removed components as described in [35]. Our aim was to achieve the best possible quantitative and qualitative description of the underlying data generating process while avoiding computational issues. All our models are detailed in *Supplementary Information: Statistical models*.

Following the standard practice for Bayesian statistics [29], we based all our model evaluations and conclusions on integrals over our models’ posterior distributions, which we calculated using adaptive Hamiltonian Monte Carlo via Stan [36] and Bambi [37]. To assess how well our models described the experimental data generating processes, we evaluated their out of sample predictive performances using expected leave-one-observation-out log predictive density. We complemented this quantitative evaluation with a qualitative assessment of agreement between our models’ posterior predictive distributions and the observed measurements based on graphical checks. The latter are shown in Supplementary Fig. 2-9 for all models used. Instructions for reproducing our analysis are available at a <u>GitHub repository</u>.

Our statistical analysis is different from null hypothesis significance testing. See [38] for a detailed description of the differences between these two methods. Our analysis does not involve null models or hypothetical unrealized datasets: The primary questions are simply whether each model adequately describes the experimental data and, if so, what *conclusion* can we make. A *conclusion* in our analysis is roughly equivalent to an output of a single hypothesis test in the frequentist analysis.

### Statistics: Test statistic and Conclusions

When we were satisfied with a model, we extracted *conclusions* from it by specifying a *test statistic* in terms of the model’s parameters and examining the marginal posterior distribution of that statistic. Here we illustrate making a conclusion with one of the results from this work: Comparing correlation coefficients, CC (quantifies the myogenic response), in adult and old mice (Fig. 3d). First, we fitted our model to the CC estimated from experimental data, yielding expected CC (i.e., the true values, unknown to us, predicted by the model) in adult, CC_Adult_, and old mice, CC_Old_.

Second, we defined a test statistic, TS = CC_Adult_ - CC_Old_, and numerically constructed its probability distribution using Monte Carlo simulation. Note how in Fig. 3d we used a simplified notation “*Adult – Old”* for this TS. We used this difference notation for all statistical analyses. Also note how we normalized all TSs so we can visualize several of them on a single plot.

Third, we visualized the TS distribution as a mean and a [2.5%, 97.5%] inter-quantile range, shown as a green point and a horizontal line, respectively, in Fig.3d. The mean of the TS (*Adult – Old*) is negative and the inter-quantile range does not include zero, so we made a *conclusion*: Aging is associated with increased CC (lower myogenic response). We can further quantify the “strength” of the effect, characterizing our certainty in the sign, “+” or “-”, of the TS, by calculating the total posterior probability mass of all TS values below zero (or above zero for a positive effect). This summed probability, we denoted as *Sign Probability* (SP). For this case, SP = 0.99. Because SP > 0.975, we concluded an effect (we use * to denote SP > 0.975 in figures). Figure 3d also visualizes another *conclusion*: AHC 2 treatment is associated with a higher CC than AHC 1. This follows from the distribution of the TS (*AHC 2 – AHC 1*) taking only positive values (SP = 1.0). The protocol outlined here was applied to all other statistical comparisons reported in this paper.

## Results

### Acute hypertensive challenge disrupts neurovascular coupling and myogenic response in the microvasculature

To study how PSs regulate the microvascular blood flow in adult and old mice during a BP surge, we followed changes in blood flow downstream of PSs in response to two consequent acute hypertensive challenges (AHC 1 and AHC 2, Fig. 1b), induced by intravenous injection of angiotensin II. Both AHCs increased mean arterial blood pressure (MAP) in adult and old mice (Fig. 1d, Table 1, Supplementary Fig. 2a).

Stimulation-induced dilation of capillaries, the neurovascular coupling (NVC), is important for delivery of energy substrates to active neurons [2,3,17,39–41]. We studied the NVC by measuring the dilation of microvessels in the whisker barrel cortex after IO stimulation. The stimulation dilated all vessel types examined, with the PS having the largest dilation and the bulb the smallest (Fig. 2a-b, Supplementary Fig. 3a, Video 1), consistent with our previous reports [1,3]. NVC responses decreased over a time course of minutes after the end of AHC 1 (Fig. 2c-d. Video 1). For example, the NVC response of 1^st^-order capillaries decreased from 19.1 ± 2.7% to 8.0 ± 1.5%, a relative decrease of 58% (averaged over adult and old mice). We did not observe any differences in NVC between adult and old mice (Fig. 2d).

**Fig. 2.**
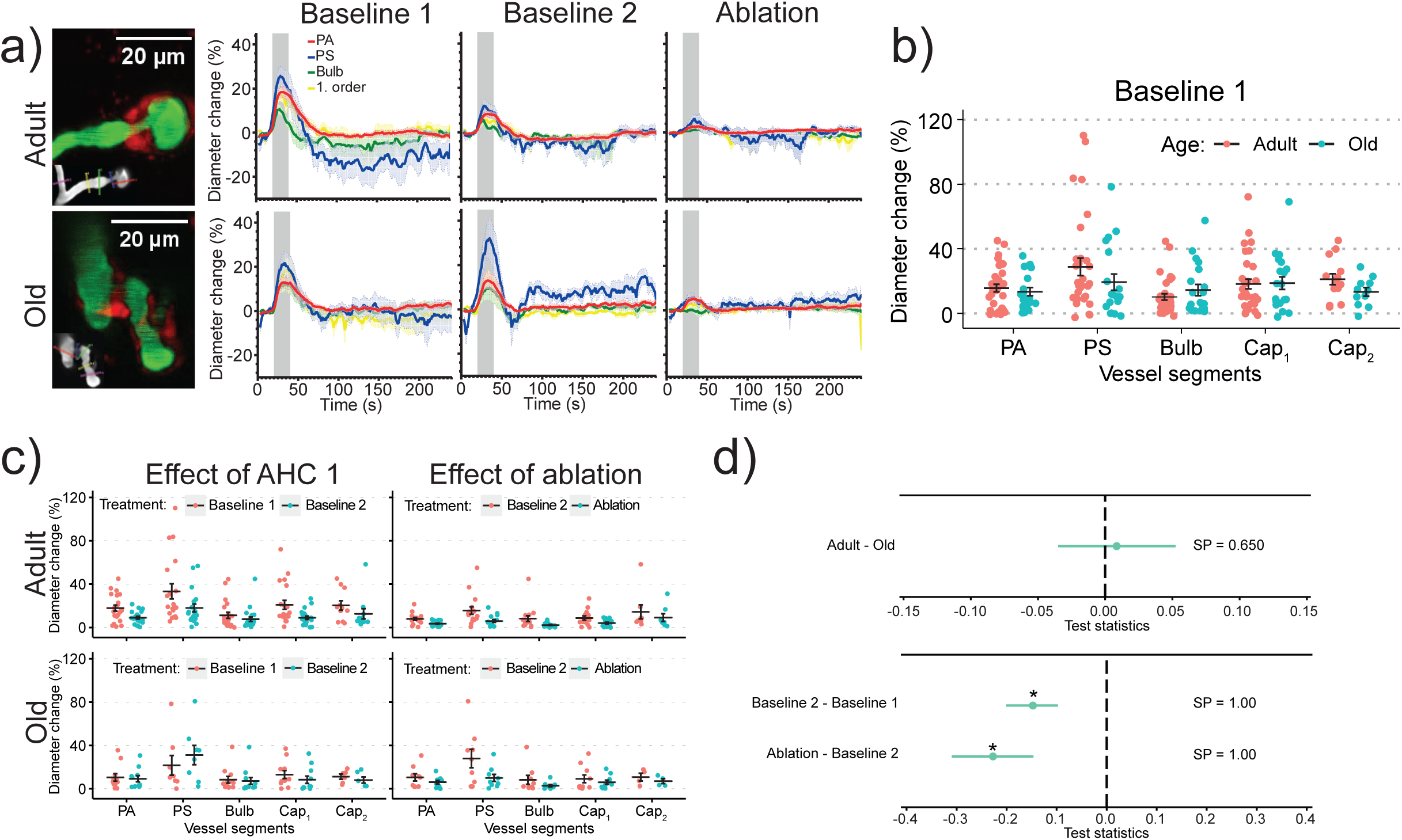
Neurovascular coupling (NVC) in adult and old mice during acute hypertensive challenge: **a**) Vessel dilations triggered by the infraorbital (IO) stimulation. **Images:** Maximum intensity projections of a volume stack of images, taken at different depths, of pericytes (NG2-dsRed, red) and blood vessels (FITC-dextran, green) around a PS. **Insets** (grey): Colored lines mark the locations of diameter measurements at PA, PS, bulb and 1-2^nd^ order capillaries. **Graphs:** Averages of traces, recorded in different mice, of relative diameter changes during IO stimulation (gray area), with shaded regions around traces showing ± s.e.m. **b)** Peak diameter changes (one point = one mouse) during IO-stimulation in adult and old mice during Baseline 1. Lines ± error bars show mean ± s.e.m. for each vessel type. **c)** Peak diameter changes (one point = one mouse) during IO-stimulation in adult and old mice after AHC 1 (left) and PS ablation (right). Note that we did not record IO-stimulated responses during AHC 1: We studied the effects of AHC 1 on NVC by comparing Baseline 2 (after AHC 1) to Baseline 1 (before AHC 1). Lines ± error bars show mean ± s.e.m. for each vessel type. **d)** Bayesian multilevel analysis of age (top) and treatment (bottom) effects on NVC. Dots and green lines show posterior mean and posterior 2.5% and 97.5% quartiles, respectively. * denotes a *conclusion*, when SP>0.975. See *Statistics: Test statistic and Conclusions* in *Methods* for more details

**Fig. 3.**
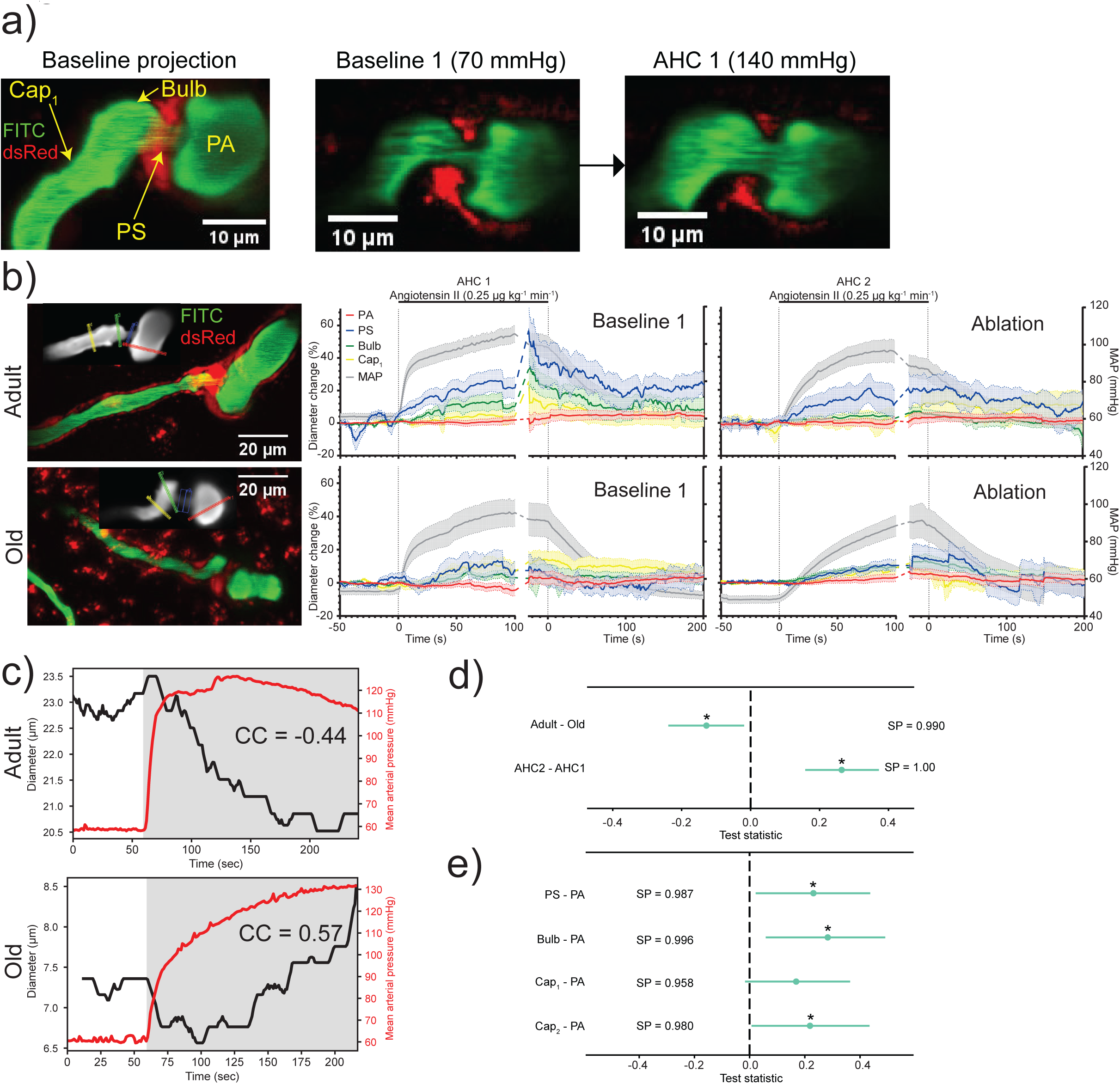
Myogenic response during acute hypertensive challenge in adult and old mice: **a**) Left: Maximum intensity projections of a volume stack of images, taken at different depths, of pericytes (NG2-dsRed, red) and blood vessels (FITC-dextran, green) around a PS. Images of the PS before (middle image) and after AHC 1 (right image) show dilations of the PS, bulb, and 1^st^-order capillary. **b) Images:** Maximum intensity projections of the volume stacks of images, taken at different depths, of pericytes (NG2-dsRed, red) and blood vessels (FITC-dextran, green), containing PSs and the surrounding vessels in adult and old mice. **Insets** (grey): colored lines mark the locations of diameter measurements for PA, PS, bulb and 1-2^nd^ order capillaries. **Graphs:** Mean MAP, averaged over different mice, and relative diameter changes during AHC 1 & 2 in adult and old mice. We aligned MAP traces as described in Fig.1d. **c)** Correlation coefficient (CC) between diameter of a PA (black) and MAP (red) estimated from the time traces in the shaded grey region in an adult (top) and an old mouse (bottom). **d-e)** Bayesian multilevel analysis of age and treatment effects (**d**) and of vessel-type effects (**e**) on CC. For panels (**d-e**), dots and green lines show posterior mean and posterior 2.5% and 97.5% quartiles, respectively. * denotes a *conclusion*, when SP>0.975. See *Statistics: Test statistic and Conclusions* in *Methods* for more details.

**Fig. 4.**
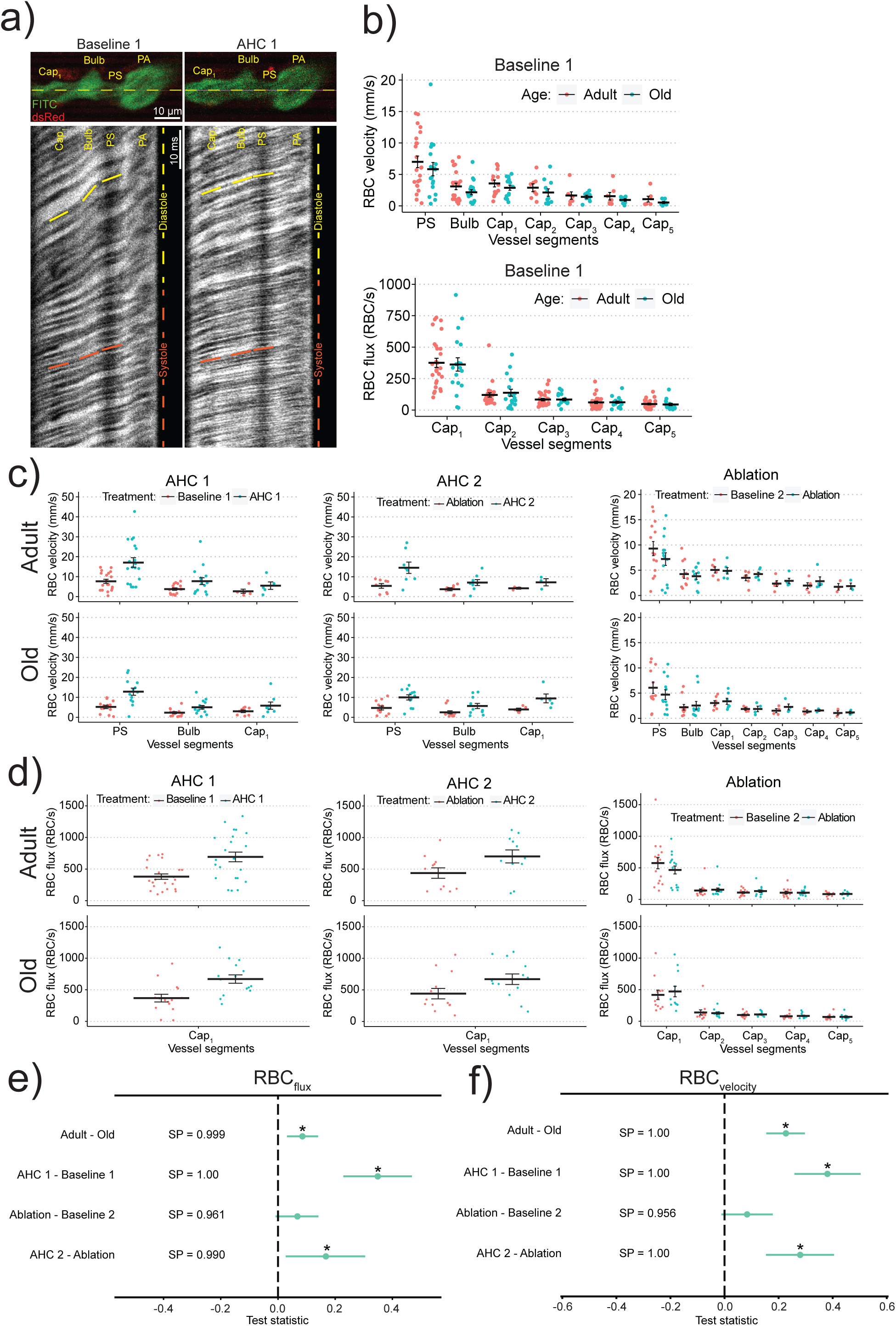
Analysis of Red Blood Cell Velocity (RBC_v_) and Flux (RBC_flux_): **a**) *Top row:* Images of pericytes (dsRed, red) and blood plasma (FITC-dextran) before and during AHC. Yellow line shows the laser beam trajectory during line-scan recording. *Lower panels:* Kymograms – line-scans of fluorescence stacked vertically in time (time increases from top to bottom) – showing RBCs as dark shadows moving from the PA to the 1^st^-order capillary (right to left). The slope of the shadows is inversely proportional to RBC_v_: Note higher RBC_v_ during the systole (red lines) compared to diastole (yellow lines), as well as higher RBC_v_ during AHC (right panels) compared to baseline (left panel). The number of RBC shadows over time estimates RBC_flux_. **b)** RBC_v_ (*top*) and RBC_flux_ (*bottom*) during *Baseline 1* in different vessel segments in adult and old mice (one point = one mouse). **c-d)** RBC_v_ (**c**) and RBC_flux_ (**d**) dependence on vessel types, treatments, and age. For panels (b-d), lines ± error bars show mean ± s.e.m. for each vessel type. **e-f)** Bayesian multilevel analysis of age and treatment effects on RBC_flux_ (**e**) and RBC_v_ (**f)**. Note how different treatments are compared to their corresponding controls: AHC 1 is compared to Baseline 1, Ablation to Baseline 2, and AHC 2 to Ablation. For panels (**e** and **f**), dots and green lines show posterior mean and posterior 2.5% and 97.5% quartiles, respectively. * denotes a *conclusion*, when SP>0.975. See *Statistics: Test statistic and Conclusions* in *Methods* for more details.

**Fig. 5.**
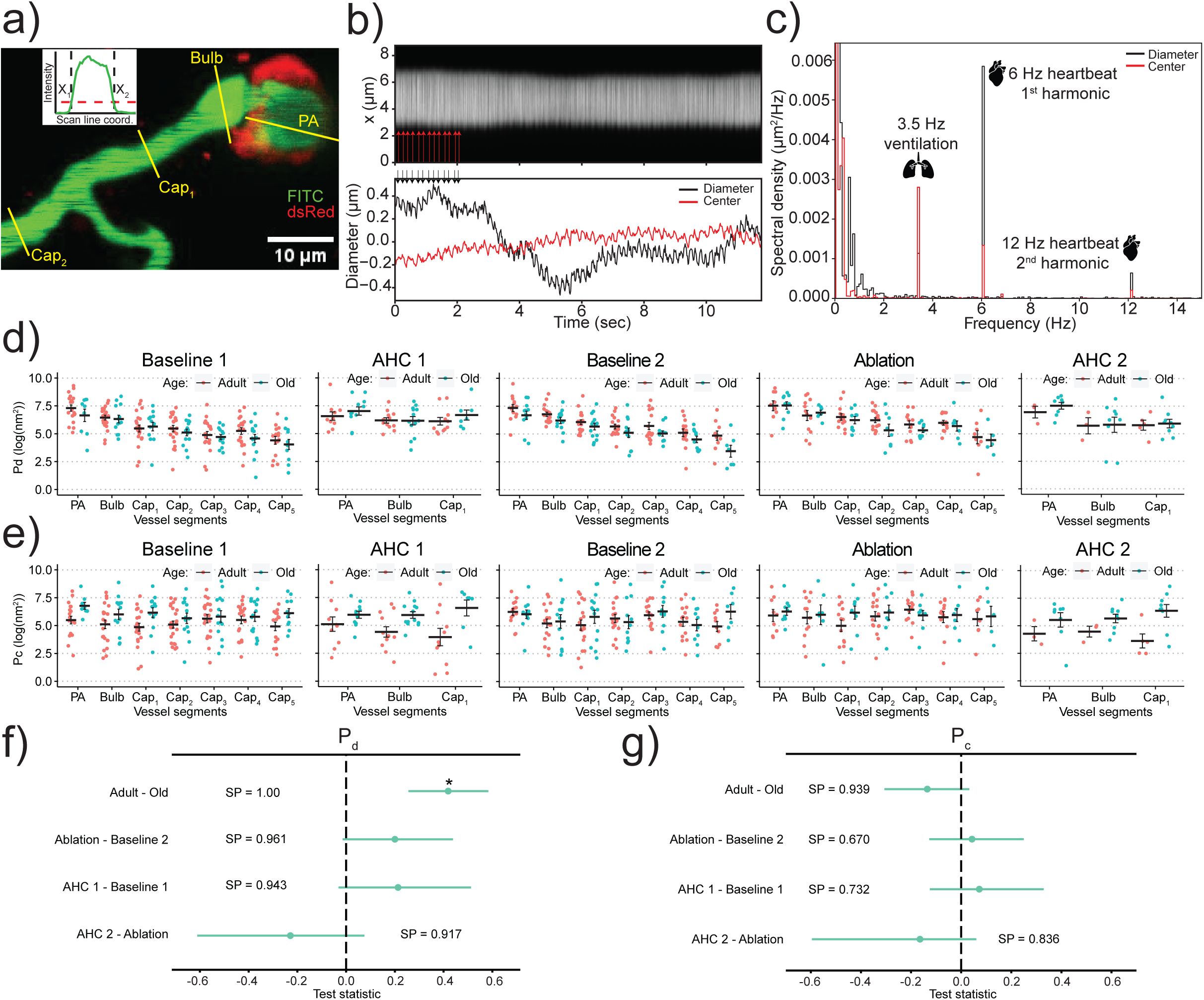
Analysis of heartbeat-driven pulsations of diameters and center positions of blood vessels: **a**) Maximum intensity projection of a volume stack of images, taken at different depths, of pericytes (NG2-dsRed, red) and blood vessels (FITC-dextran, green) around a PS. Yellow lines show trajectories of the laser beam during line-scan recordings. **Inset:** A line-distribution of FITC-dextran fluorescence (an average of raw line-scans) taken perpendicular to a vessel’s axis. The diameter of the vessel was estimated by the width of the line-distribution (see details in Methods). **b)**: Top: Kymogram of FITC-dextran fluorescence (line-scans in time) of a 1^st^-order capillary (red arrowheads indicate heartbeats). Bottom: Estimated mean-subtracted diameter (black) and center position (red) of the capillary (black arrowheads indicate heartbeats). **c)**: Power spectra of the center position (red) and diameter (black) of the corresponding time traces in (b). Organ drawings from https://scidraw.io/. **d-e)** P_d_ (d), and P_c_ (e) dependence on vessel types, treatments, and age (one point = one mouse). Lines ± error bars show mean ± s.e.m. for each vessel type. **f-g)** Bayesian multilevel analysis of age and treatment effect on P_d_ (**f**) and P_c_ (**g**). For panels (**f-g**), dots and green lines show posterior mean and posterior 2.5% and 97.5% quartiles, respectively. * denotes a *conclusion*, when SP>0.975. See *Statistics: Test statistic and Conclusions* in *Methods* for more details.

**Fig. 6.**
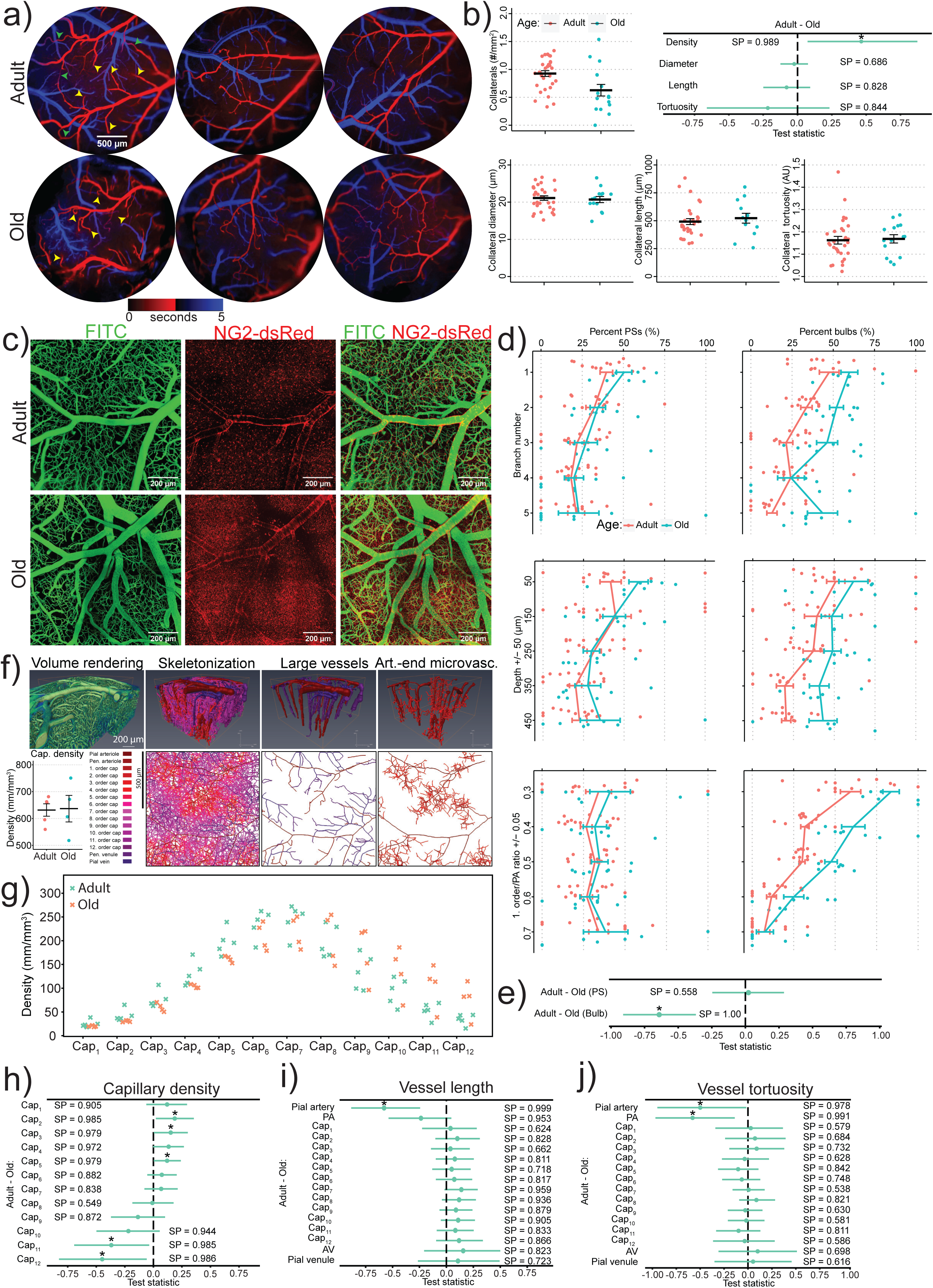
Microvascular topology of Adult and Old Mice: **a**) Three pairs (adult vs old) of diagrams, where the color of each pixel encodes the time of FITC-dextran arrival (after i.v. injection) at the pixel location: Red shows arterioles, blue – venules. Yellow and green arrowheads point to PAs and collaterals, respectively, illustrating e.g., increased PAs tortuosity in old mice. **b**) Collateral density, diameters, lengths, and tortuosity in adult and old mice. Each data point is an average over multiple measurements from a single mouse. Top right panel show Bayesian multilevel analysis of age effects on collateral density, diameter, length, and tortuosity. Dots and green lines show posterior mean and posterior 2.5% and 97.5% quartiles, respectively. * denotes a *conclusion*, when SP>0.975. **c)** Maximum-intensity projections of volume stacks of images, taken at different depths, of pericytes (NG2-dsRed, red) and blood vessels (FITC-dextran, green). The red background fluorescence in the old mice is predominantly attributed to lipofuscin [23]. **d)** Proportion of PSs and bulbs at each branchpoint, their depth, and the ratio between 1^st^-order capillary and PA’s diameters (see Methods: Quantification of precapillary sphincters and bulbs). **e)** Bayesian multilevel analysis of age effects on PS and bulb occurrence probability. Dots and green lines show posterior mean and posterior 2.5% and 97.5% quartiles, respectively. * denotes a *conclusion*, when SP>0.975. **f)** 3D reconstruction of blood vessels, accompanied by a skeletonization (lower panels) and volume representation (upper panels) of the segmented microvasculature. The lower left panel compares the overall capillary density between adult and old mice. **g)** The distribution of capillary densities in adult and old mice. **h-j)** Bayesian multilevel analysis of age effects on vessel density **(h)**, length **(i)**, and tortuosity **(j)**. For panels **(h-j)**, dots and green lines show posterior mean and posterior 2.5% and 97.5% quartiles, respectively. * denotes a *conclusion*, when SP>0.975. See *Statistics: Test statistic and Conclusions* in *Methods* for more details.

During peak AHC, we observed dilation of PSs and downstream capillaries (Fig. 3a-b, Video 2), indicating an impaired myogenic response. To quantify the microvascular myogenic response to AHC, we estimated the Pearson correlation coefficient (CC) – a commonly used metric to study autoregulation [42,43] – between the time series of MAP and the vessel’s diameter (Fig. 3c). A positive CC indicates an impaired myogenic response (a vessel dilates passively, driven by increasing pressure), while a negative CC indicates functional myogenic response (a vessel constricts in response to increasing pressure). Average CCs were positive for both adult and old mice, indicating a compromised myogenic response during AHC. Old mice showed a higher overall (both AHC 1 and 2) CC compared to adult mice (Fig. 3d, Supplementary Fig. 4), suggesting a weaker myogenic response in old mice. The second AHC further increased CC compared to the first AHC (Fig. 3d), indicating a further decreased myogenic response, which may be caused by a dysfunctional PS or by an accumulated damage to the microvessels caused by the two consequent AHC. Vessels downstream PA showed a higher CC than in the PA, except for the 1^st^ order capillary (Fig. 3e), illustrating how myogenic responses vary along the microvascular tree.

Adult mice showed higher RBC_v_ and RBC_flux_ compared to old mice (Fig. 4a-b, e-f, Supplementary Fig. 5). Both AHC 1 and AHC 2 increased both RBC_v_ and RBC_flux_ (Fig. 4c-f). For example, during the first and second AHC, the 1^st^-order capillary RBC_v_ increased from 3.2 ± 0.4 mm/s to 5.8 ± 1.3 mm/s (81% increase) and from 4.1 ± 0.5 mm/s to 8.7 ± 1.5 mm/s (121% increase), respectively (averaged over both adult and old mice).

### Heartbeat-induced diameter pulsations of microvessels are decreased in old mice

Heartbeats produce two pressure waves in the brain. One wave propagates along the vasculature and dilates the vessels at the heartbeat frequency. The second wave propagates through the brain tissue and cerebrospinal fluid [44], driven by pulsations of large brain arteries, and causes the brain tissue to oscillate at the heartbeat frequency [45]. We quantified the effect of the first and second waves on the vasculature by estimating spectral power of the first harmonic of the heartbeat pulsations of a vessel’s diameter, P_d_, and its center position, P_c_ (Fig. 5a-c). P_d_ and P_c_ are proportional to the squared amplitudes of vessels diameters and center position pulsations, respectively. A reduced P_d_ may indicate a reduced PP, which drives the pulsations of the vessel wall, or an increased stiffness of the vessel wall. Consistent with the decaying pressure wave propagating from the arterial to the venous end of the circulation (and assuming the same stiffness of different studied vessels), P_d_ gradually decreased from PAs to 5^th^-order capillaries (Fig. 5d, Supplementary Fig. 6a). In contrast, P_c_ was the same for all vessels (Fig. 5e, Supplementary Fig. 6a), because the centers of all vessels moved together with the brain volume containing them, which moved due to the pressure wave propagating through the brain tissue.

P_d_, but not P_c_, were lower in old mice compared to adult mice (Fig. 5f-g), which can be explained by increased stiffness (i.e. higher Young’s modulus) of microvessels in the old mice. Note that a lower P_d_ of the brain microvasculature cannot be explained by a difference in systemic PP, which was similar between adult and old mice (Table 1, Supplementary Fig. 2a). Also, the lower P_d_ in old mice cannot be explained by an increased baseline diameter (which may result in an increased stiffness of the vessels wall; hence, decreased P_d_). On the contrary, old mice showed lower diameters of 1^st^ order capillaries, and no differences in diameters for other vessel types compared to the adult mice (Supplementary Fig. 7a). Finally, unchanged P_c_ suggests that the amplitude of pulsations of large brain arteries, creating pressure waves in the brain tissue, did not change. Given unchanged PP, this result may indicate a similar stiffness of the large brain arteries in adult and old mice (assuming similar stiffness of the brain tissue in adult and old mice). Taken together, we explain lower P_d_ in old mice by the increased stiffness of the microvasculature.

### Damage to precapillary sphincters disrupts neurovascular coupling, does not affect downstream blood flow, and does not change the amplitudes of pulsations of capillary diameters

PS ablation increased the mean vessel lumen diameter at the site of the PS on average, from 4.01±0.34 µm to 5.31±0.25 µm (32% increase) for adult and from 4.54±0.55 µm to 5.91±0.71 µm (30% increase) for old mice (Fig. 1c). Following the PS ablation, NVC responses decreased (Fig. 2d, bottom), for example from 8.0 ± 1.5% to 3.9 ± 0.8% for 1^st^-order capillaries, a relative decrease of more than 50% (averaged over both adult and old mice). We did not find an effect of PS ablation on RBC_v_ and RBC_flux_ (Fig. 4c-f), or an effect of PS ablation on P_d_ and P_c_ in the downstream capillaries (Fig. 5f-g).

### Brain microvascular topology changes with aging

To better understand the age-related differences in vascular functions, we studied the topology of the vasculature of adult and old mice. We observed the following changes in microvascular topology of old mice, compared to the adult: (i) reduced number of pial collaterals (Fig. 6a-b, Supplementary Fig. 8); (ii) decreased capillary density in the arteriolar end and increased in the venous end (Fig. 6g-h, Supplementary Fig. 9); (iii) increased length and tortuosity of pial arterioles and PAs (Fig. 6i-j, Supplementary Fig. 9); (iv) increased number of bulbs at arteriolar branchpoints (Fig. 6d-e). The largest differences in bulb percentage were found in cortical layers 3 and 4 (350±50 µm depth, adult layer 3: 19.7±5.2%, old layer 3: 40.8±6.0%) and for relatively larger PAs compared with the 1^st^-order capillaries (Fig. 6d, 1^st^-order capillary/PA diameter ratio = 0.3: adult = 59.3±3.7%, old = 83.7±6.0%).

## Discussion

### Myogenic response is overpowered by acute hypertensive challenge and impaired by precapillary sphincter ablation

A rapid BP increase (up to 80% within minutes) compromised the microvascular myogenic response, causing capillary dilation, shown by a positive CC between MAP and microvascular diameter. The higher CC in old mice indicated a more compromised response compared to adults. This impairment may trigger blood-brain barrier disruption, linked to age-related cerebrovascular pathologies [10,17,46]. The increased CC after PS ablation suggests PSs may help execute a myogenic response, protecting the capillary bed from abrupt BP surges.

### Neurovascular coupling is disrupted by acute hypertensive challenge and by precapillary sphincter ablation

AHC reduced NVC response, possibly due to inhibitory effects of angiotensin II [40,47]. PS ablation also reduced NVC response. PSs are located at the junctions between PAs and the 1^st^-order capillaries, which are hubs for myoendothelial junctions [4,5] – structures that facilitate NVC by providing communication between endothelial cells and pericytes/smooth muscle cells [48,49].

Thus, ablation of PSs, by damaging the first order capillaries, can potentially disrupt the NVC. Finally, unlike our previous study [3], we found no substantial difference in NVC responses between adult and old mice around PSs, possibly due to fewer replicates [3].

### Microvascular blood flow and vessel diameter pulsations decrease in old mice and are not affected by precapillary sphincter ablation

We found that old mice have decreased RBC_flux_, RBC_v_, and P_d_. The latter could be explained by stiffer walls of microvasculature of old mice. The decrease in blood flow can be partly explained by the decreased MAP in old mice. We hypothesised that PS ablation could lead to an increase in RBC_flux_, RBC_v_, and P_d_ in the downstream capillaries. The three quantities, however, remained unchanged following the PS ablation. One can speculate that upstream or downstream vessels may actively compensate the ablation-induced dilation of the PSs to keep the blood flow and pulsatile stress at a constant level. That way, if one of the vessels, connected in a series, fails to sustain the blood flow, the other vessels help stabilise it. On the other hand, while the results do not satisfy our test criterion of at least 97.5% positive (or negative) posterior mass (see *Statistics: Test statistic and Conclusions* in *Methods*), there is a clear trend in favour of an increase of RBC_flux_ (SP=0.96, Fig. 4e), RBC_v_ (SP=0.96, Fig. 4f), and P_d_ (SP=0.96, Fig. 5f) after PS ablation. Thus, more research is needed to determine whether PSs can help absorb heartbeat-induced pulsations in brain microvessels, similarly to how large arteries absorb pulsations caused by the beating heart [50,51].

### Old mice lose capillaries near the arterial end and gain capillaries near the venous end

Age-associated remodeling in arterial-end microvascular topology, including increased arteriole tortuosity, reduced collaterals, and reduced capillary density, agrees with previous studies [3,18–20,52–55]. Reduced cerebral capillary density is the main factor explaining decreased blood flow in aging rats [56]. We observed loss of arterial-end capillaries and gain of venous-end capillaries in old mice, likely due to arterial-end capillary plugging and venous-end compensatory angiogenesis [18–20,55,57]. The impact of this remodeling on microvascular resistance is unclear due to the lack of experimental studies.

### Old mice have more bulbs

We found more bulbs (distensions of 1^st^-order capillaries downstream of PSs) in old mice, especially where the first-order capillary diameter is less than half the upstream PA diameter. In this case, a small increase in PS diameter may induce a large increase in pressure downstream, where the bulb is located. For example, according to theoretical simulations, a five-fold dilation of a PS from 20% to 100% of the diameter of the downstream first-order capillary, increases the pressure downstream of the PS by 20-30 times (see Fig. 8 in [58]). Bulbs lack ensheathing pericytes [1,4], making their vessel walls more compliant and prone to damage from surging blood pressures. We suggest that bulbs are capillary aneurysms developing over a mouse’s lifetime due to repeated blood pressure surges and irreversible stretching of capillary walls downstream of the PS.

## Summary

Our findings highlight the important role of PSs in sustaining the neurovascular coupling and, possibly, the myogenic response, under elevated BP. A large and brief systemic BP increase (∼70% for 8 min), induced by angiotensin II, disrupted the myogenic response and NVC, and increased cerebral blood flow in both adult and old mice. Old mice showed decreased pulsations of microvessel diameters, possibly due to increased stiffness of their walls compared to adult mice.

Finally, we observed ageing-related microvascular remodeling, including the loss of arterial-end capillaries, gain of venous-end capillaries, and increased aneurysm-like capillary distensions, bulbs, downstream of PSs. These microvascular changes may be adaptations to detrimental events during brain aging but could also affect blood flow.

## Acknowledgements

We thank Micael Lønstrup for training and assistance with microsurgery. We thank Dr. Changsi Cai for her Matlab code for estimating diameters of blood vessels from 3D image stacks.

## Supplementary video captions

**Video 1:** Example of IO stimulation before/after AHC and a PS ablation in an adult and an old mouse.

**Video 2:** AHCs dilate PSs and microvasculature.

**Video 3:** Leakage of FITC to the perivascular space recorded after AHC 2.

**Video 4:** Cortical angiograms of an adult and an old NG2-DsRed mouse, where the color encodes the relative arrival time, i.e. red – early (arterioles) and blue – late (veins), of FITC-dextran fluorescence after its i.v. injection.

**Video 5:** Combining multiple volume-image-stacks into a single map of the cortical microvascular topology recorded in an adult and an old mouse *in vivo*. We skeletonized and labeled each vascular bifurcation using Amira.

