## Supplementary information for "Brain precapillary sphincters modulate myogenic tone in adult and aged mice"

### 1 Supplementary figures

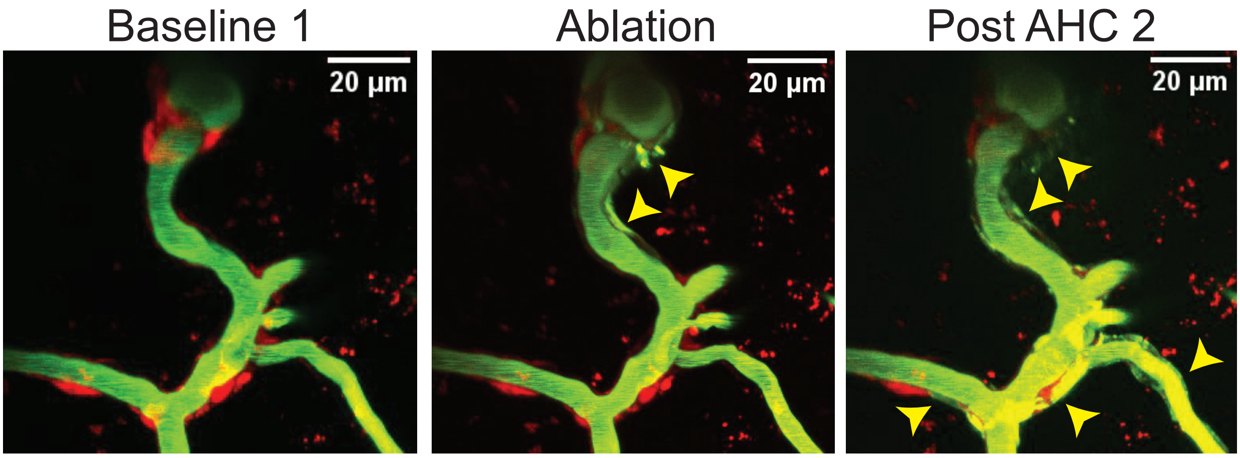

Supplementary Fig. 1: Images of blood vessels with FITC-dextran in blood plasma (green) and DsRed in pericytes and smooth muscle cells (red), showing leakage of FITC-dextran (yellow arroheads) near a PS after its ablation. Left image shows the vasculature before any treatment, middle image — after PS's ablation, and right image — after AHC 2. Note how FITC-dextran propagated along the vasculature, possibly in the perivascular spaces.

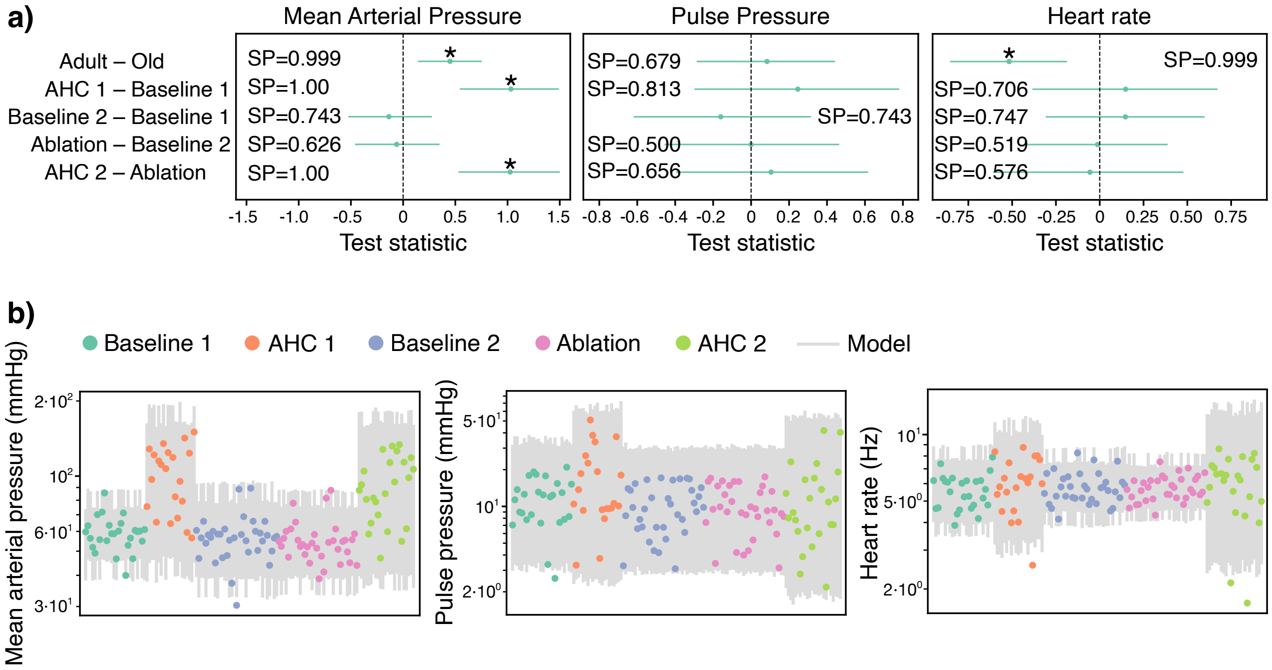

Supplementary Fig. 2: Analysis of mean arterial pressure (MAP), pulse pressure (PP), and heart rate (HR). For the definition of the models see section 2.5. **(a)** Age and treatment effects on MAP (left), PP (middle), and HR (right). Dots and green lines show posterior mean and posterior 2.5% and 97.5% quartiles, respectively. AHC1 and AHC2 were associated with increased MAP but not PP and not HR. Aging was associated with decreased MAP and increased HR. * denotes a *conclusion*, when SP$>$ 0.975. See *Statistics: Test statistic and Conclusions* in Methods for more details. **(b)** Measurements of MAP (left), PP (middle), and HR (right), shown as dots, plotted alongside the corresponding statistical model's 1%-99% marginal posterior predictive intervals. Measurements are sorted by treatment but otherwise in random order. The plots show that the three models fit MAP, PP, and HR adequately, with most measurements contained within the corresponding marginal interval and no obvious pattern in the deviations.

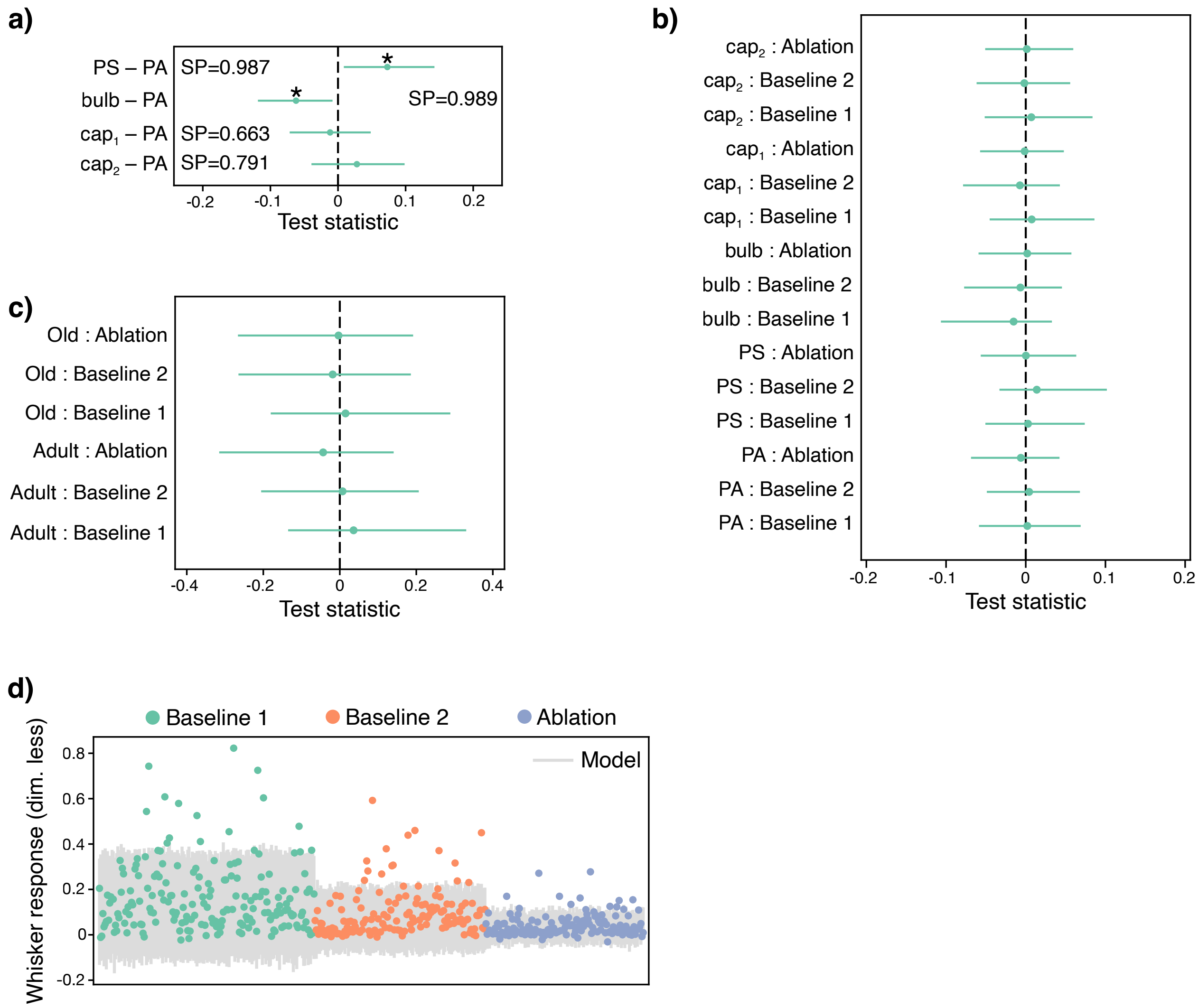

Supplementary Fig. 3: Analysis of whisker responses (WR). For the definition of the models see section 2.6. **(a)** Vessel-type effects on WR: PS and bulbs are associated with highest and lowest WRs, respectively. Dots and green lines show posterior mean and posterior 2.5% and 97.5% quartiles, respectively. * denotes a *conclusion*, when SP$>$ 0.975. See *Statistics: Test statistic and Conclusions* in Methods for more details. **(b)** Interaction effects between vessel-types and treatments. All interaction effects were consistent with zero (SP$<$ 0.975). **(c)** Interaction effects between age and treatments. All interaction effects were consistent with zero (SP$<$ 0.975). **(d)** Measurements of WR, shown as dots, plotted alongside the model's 1%-99% marginal posterior predictive intervals. Measurements are sorted by treatment but otherwise in random order. Unlike for other models, there is a systematic pattern in the model's bad predictions: The model was not able to explain the highest WRs in all three treatment categories. These measurements were probably caused by a different process than the other measurements: In order to identify and quantitatively characterize these outliers one can use a mixture model with two likelihoods—one for each postulated process—as well as a mixing probability, indicating which process has the best chance of having generated any measurement. This more complex model would potentially be a more accurate description of our experiment. However, since there were relatively few extreme outliers (10 out of 475 measurements were above the 99.9th posterior predictive percentile), we judged that extending our model in this way would have a small impact on our main results, and therefore decided not to do so.

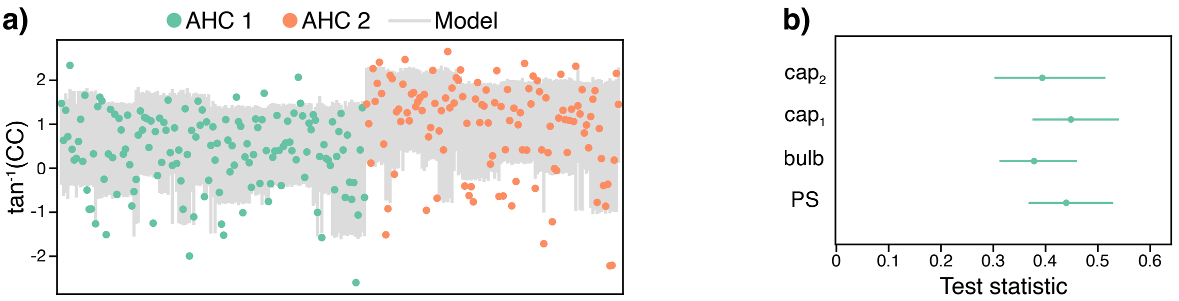

Supplementary Fig. 4: Analysis of the myogenic response, quantified as the correlation coefficient, CC, between a vessel's diameter and MAP. For the definition of the models see section 2.7. **(a)** Measurements of CC, shown as dots, plotted alongside the model's 1%-99% marginal posterior predictive intervals. Measurements are sorted by treatment but otherwise in random order. The fit to the data is reasonably good, though worse for the AHC2 treatment. Similar to the model of whisker responses (section 2.6), we judged that improving the fit by expanding our statistical model would not substantially change our results. **(b)** Marginal posterior distributions for measurement error parameters, which varied by vessel type. According to our model, measurements of bulbs and second order capillaries tended to be more accurate than measurements of sphincters and first order capillaries.

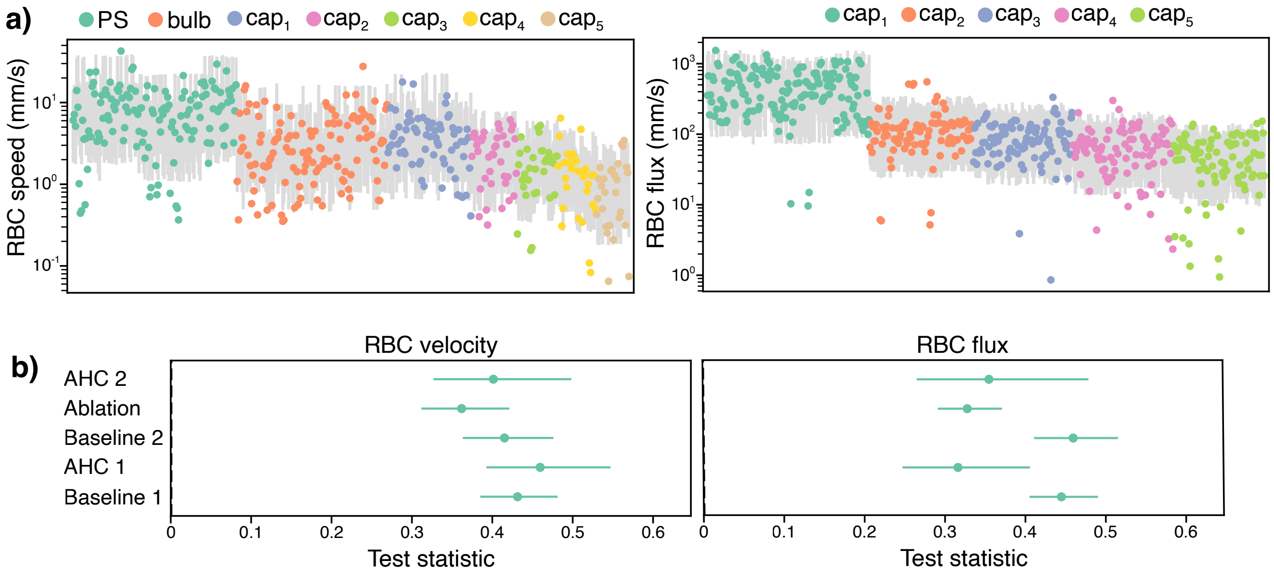

Supplementary Fig. 5: Analysis of RBCs velocity and flux. For the definition of the models see section 2.8. **(a)** Measurements of RBCs velocity (left) and RBCs flux (right), shown as dots, plotted alongside the models 1%-99% marginal posterior predictive intervals. Measurements are sorted by vessel-types but otherwise in random order. Both plots show a few outlier measurements for each vessel-type, mostly on the low side, i.e. where the measured speed or flux was much lower than our models expected. We were not able to extend our models to accommodate these outliers and, therefore, judged that they were caused by a process that we did not consider. **(b)** Marginal posterior distributions for measurement error, which varied by treatment. According to our model, measurements of RBC velocity were most accurate for the ablation treatment and least accurate for treatment AHC 1. Measurements of RBC flux were, estimated to be less accurate for the baseline treatment than for the other treatments.

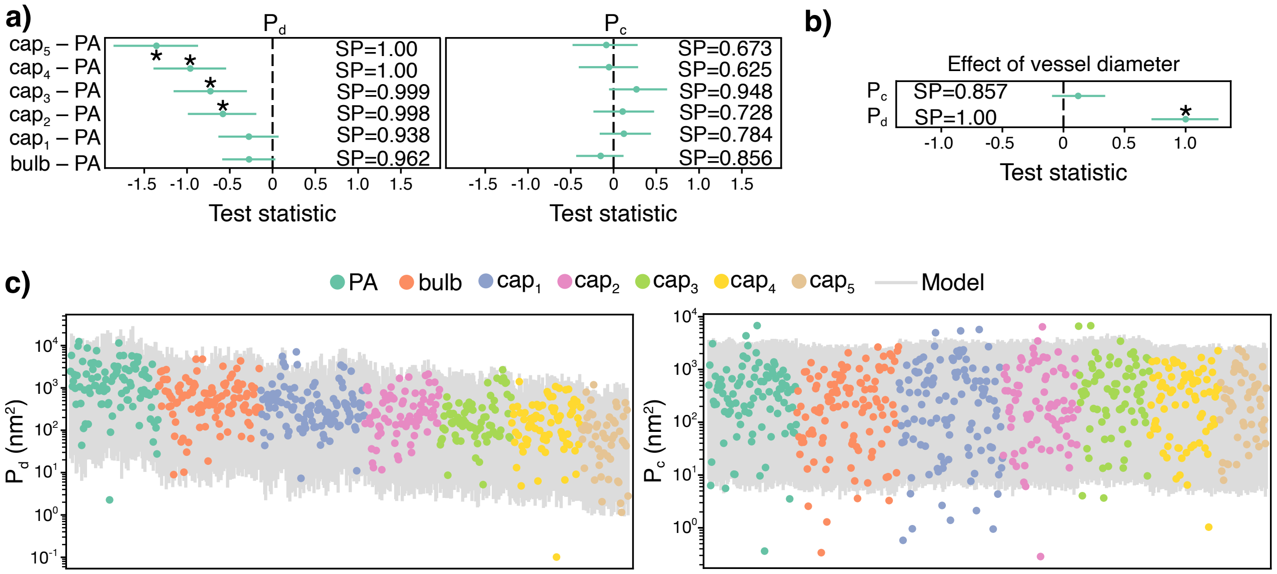

Supplementary Fig. 6: Analysis of $P_d$ and $P_c$. For the definition of the models see section 2.9. **(a)** Vessel-type effects on $P_d$ (left) and $P_c$ (right): Increased capillary order is associated with decreased $P_d$ but not $P_c$. **(b)** In panels **a,b** * denotes a *conclusion*, when SP> 0.975. See *Statistics: Test statistic and Conclusions* in Methods for more details. Increased vessel diameter is associated with higher $P_d$. There is no effect of a vessel's diameter on $P_c$. **(c)** Measurements of $P_d$ (left) and $P_c$ (right), shown as dots, plotted alongside the models 1%-99% marginal posterior predictive intervals. Measurements are sorted by vessel-type but otherwise in random order. 34 out of 601 $P_c$ measurements (≈ 5.5%) are outside the plotted intervals, i.e., more than the expected value of 2%. We did not find any systematic pattern in these large errors.

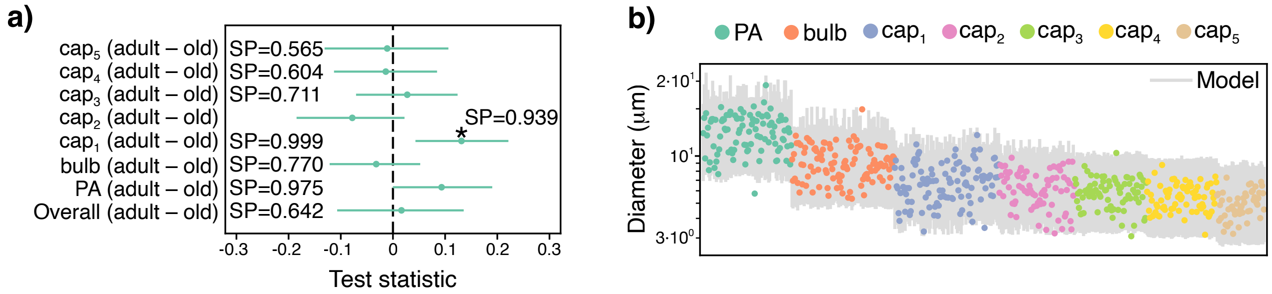

Supplementary Fig. 7: Analysis of baseline diameters. For the definition of the models see section 2.10. **(a)** Age effect on the baseline diameters of various vessel-types. Note the absence of the age effect on all vessel types combined. * denotes a *conclusion*, when SP> 0.975. See *Statistics: Test statistic and Conclusions* in Methods for more details. **(b)** Measurements of baseline diameters, shown as dots, plotted alongside the model's 1%-99% marginal posterior predictive intervals. Measurements are sorted by vessel type but otherwise in random order. There are no obvious problems with the fit to the observed data.

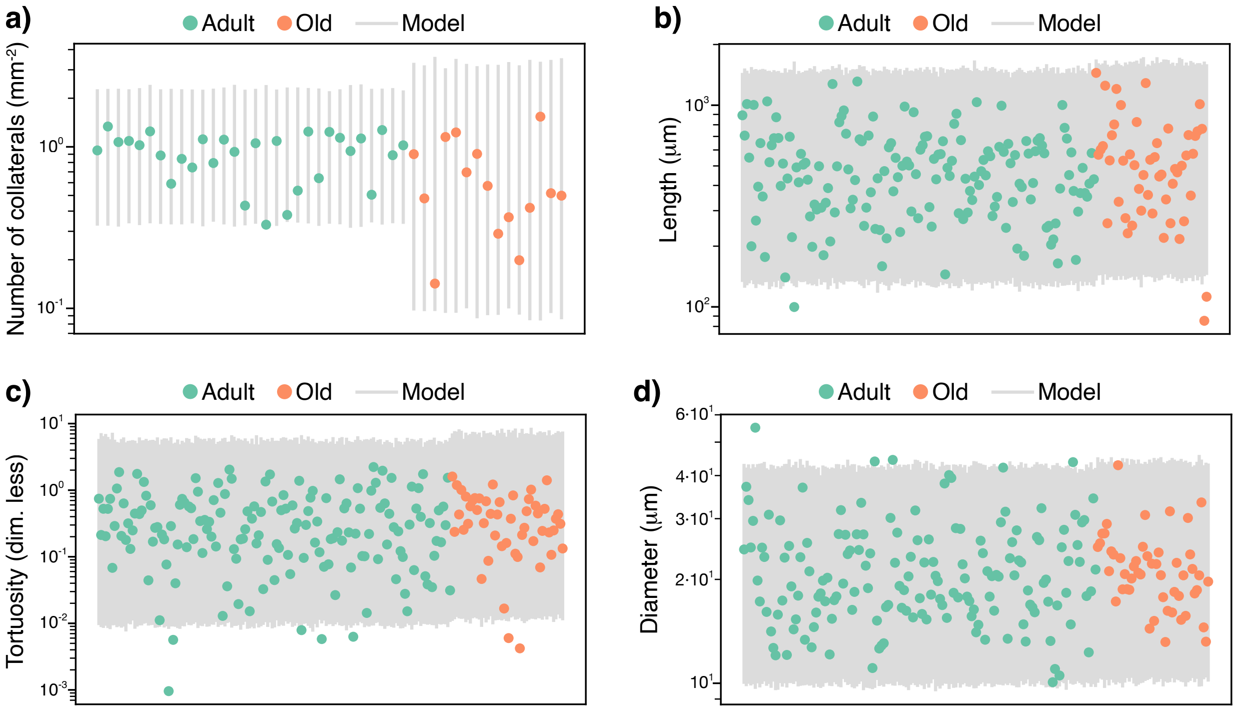

Supplementary Fig. 8: Analysis of topology of collaterals on the brain's surface. For the definition of the models see section 2.11. **(a)** Measurements of the number of collaterals, shown as dots, plotted alongside the model's 1%-99% marginal posterior predictive intervals. **(b)** Measurements of the length of vessels, shown as dots, plotted alongside the model's 1%-99% marginal posterior predictive intervals. **(c)** Measurements of the tortuosity of vessels, shown as dots, plotted alongside the model's 1%-99% marginal posterior predictive intervals. **(d)** Measurements of the diameters of vessels, shown as dots, plotted alongside the model's 1%-99% marginal posterior predictive intervals. In all four panels the measurements are sorted by age but otherwise in random order. The fit to the measurements is broadly acceptable, though, perhaps due to the fewer available covariates, the model predictions do not vary greatly for different measurements.

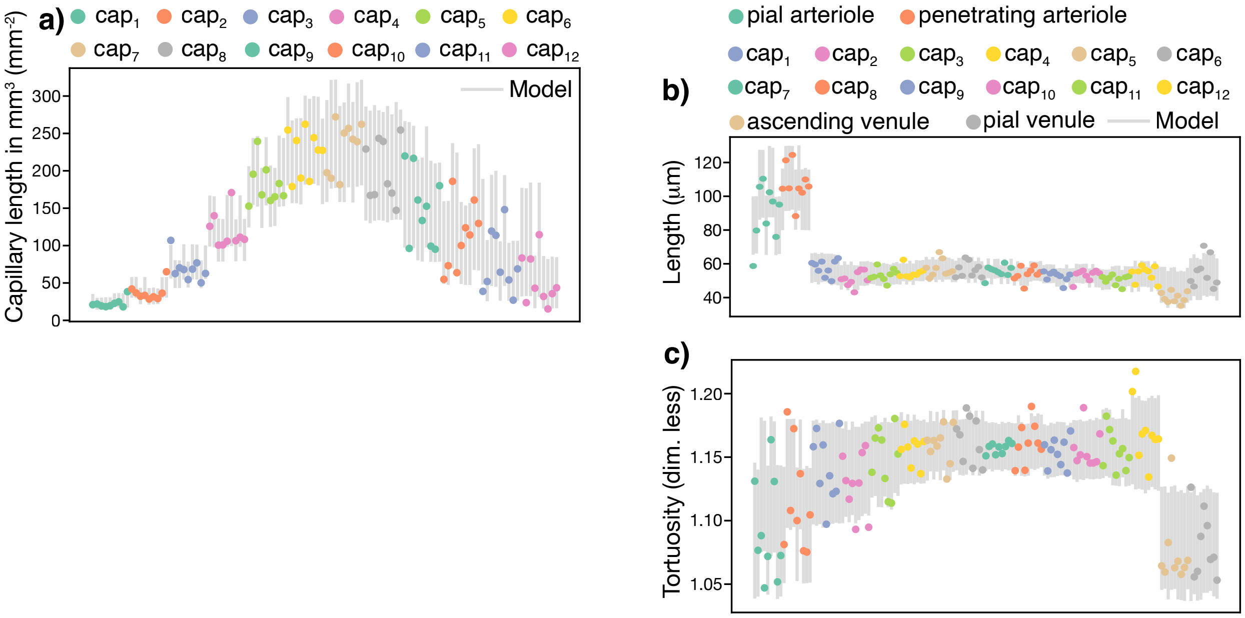

Supplementary Fig. 9: Analysis of topology of capillaries on the brain's surface. For the definition of the models see section 2.12. **(a)** Measurements of capillary density, shown as dots, plotted alongside the model's 1%-99% marginal posterior predictive intervals. **(b)** Measurements of the length of capillaries, shown as dots, plotted alongside the model's 1%-99% marginal posterior predictive intervals. **(c)** Measurements of the tortuosity of capillaries, shown as dots, plotted alongside the model's 1%-99% marginal posterior predictive intervals. In all four panels the measurements are sorted by vessel type but otherwise in random order. The fit to the measurements is broadly acceptable.

#### 2 Statistical models

Our overall statistical modelling strategy was to (i) construct Bayesian multilevel generalised linear regression models describing our data, (ii) perform posterior sampling using Markov Chain Monte Carlo, and (iii) analyse the distribution of test statistics of interest in the posterior samples.

We selected a model depending on the nature of the measurements, the structural and quantitative prior knowledge available, and the performance of our models. We followed the Bayesian Workflow described by [5] et al., starting with a simple model and iteratively adding and removing components, or adjusting priors and hyperparameters, aiming to achieve a satisfactory qualitative and quantitative description of the data generating process while avoiding computational issues. The final models used in each case are described below.

##### 2.1 Reproducibility

Instructions for reproducing our analysis are available at https://github.com/teddygroves/sphincter/blob/main/README.md.

##### 2.2 Bayesian posterior sampling

Our main results describe the distribution across posterior samples of test statistics, i.e. deterministic functions of our models' parameters, e.g., the difference in treatment effects. We summarized the distributions of these test statistics with the mean and 2.5% to 97.5% inter-quantile ranges. These ranges were chosen to indicate the range of values that are consistent with the assumptions that the model encodes about the experimental setup. For example, where we report the 1% quantile for a test statistic, this means that 99% of posterior samples of the statistic were greater than this value, and can be interpreted as indicating that the model's probability of a lower value is approximately 1% (allowing for a small Monte Carlo error, typically less than 1% of the posterior mean). As with any statistical analysis, the interpretability and meaningfulness of the results depend on the model accurately approximating the true data generating process and on the correctness of the computational method we used to obtain posterior samples.

##### 2.3 Model assessment and validation

We assessed models quantitatively using the estimated leave-one-observation-out log predictive density score [8], and qualitatively using graphical prior and posterior predictive checks such as those presented in this document.

We validated our statistical computation using standard Hamiltonian Monte Carlo diagnostics, including the improved $\hat{R}$ statistic [9] as well as inspection for post-warmup divergent transitions [2], problematic EBFI statistics, tree depth or effective sample size to total sample size ratios. All reported models had improved $\hat{R} \approx 1$ and no divergent transitions or other signs of algorithm failure.

##### 2.4 The base model

We used generalised linear regression models to describe the information in our measurements. We modelled the expected value of any measurement $y$, given measurement distribution $p$ as $E_p(y|x, \beta) = f^{-1}(\eta)$, where $\eta$ is a linear predictor obtained by applying linear operations to some regression coefficients $\beta$ and covariates $x$, and $f$ is a link function connecting $\eta$ with the possibly non-linear range of possible values of $y$. The link function, regression coefficients and measurement distribution varied from case to case, as described below. To capture structural knowledge about the experimental setup, for example potential similarities between measurements of mice in the same treatment or vessel type category, we used hierarchical parameters. Our *base model* ($\eta_{atv}$) on which most of our models are based on, can be written as

$$
\begin{aligned}
\eta_{atv} &= \mu_a^{\text{age}} + \alpha_t^{\text{treatment}} + \alpha_v^{\text{vessel type}} \\
\alpha^{\text{treatment}} &\sim N(0, \tau^{\text{treatment}}) \\
\alpha^{\text{vessel type}} &\sim N(0, \tau^{\text{vessel type}}) \\
\mu^{\text{age}} &\sim N(a1, s1) \\
\tau^{\text{treatment}} &\sim N^{+}(0, s2) \\
\tau^{\text{vessel type}} &\sim N^{+}(0, s3)
\end{aligned} \tag{1}
$$

$N$ is the normal distribution and $N^+$ is the half-normal distribution, i.e. normal distribution with support only for non-negative numbers. $a1$, $s1$, $s2$ and $s3$ are user-provided hyperparameters.

This structure is hierarchical because the prior distributions for parameters $\alpha^{\text{treatment}}$ and $\alpha^{\text{vessel type}}$ depend on other parameters, specifically $\tau^{\text{treatment}}$ and $\tau^{\text{vessel type}}$. The $\tau$ parameters function to control the general dispersion away from zero of their corresponding $\alpha$ parameters, allowing the model to infer both the general prevalence of treatment and vessel type effects and the values of any specific effects.

Below we describe the distinctive mathematical features of each model that we used in our final analysis. Code implementing all of the full models, as used, is available in the code repository and can be inspected to clarify implementation details or to find the values of model settings not mentioned below, such as the values of user-specified prior parameters. All analyses except for the collateral and branchpoint analyses have corresponding Stan files in the folder `https://github.com/teddygroves/sphincter/tree/main/sphincter/stan`. For the collateral and branchpoint analyses we used simpler models implemented with PyMC via the library bambi, which are available at `https://github.com/teddygroves/sphincter/blob/main/sphincter/collaterals.py` and `https://github.com/teddygroves/sphincter/blob/main/sphincter/branchpoints.py`. Note that in some cases we relied on bambi's default prior configurations rather than setting priors explicitly.

#### 2.5 Blood pressure and heart rate

We modeled the measurements of mean arterial pressure (MAP), pulse pressure (PP), and heart rate (HR) on log scale, using three different models (Supplementary Fig. 2). Our final model includes varying measurement error parameters to accommodate the heteroskedastic data and effects for age, treatment and age:treatment interaction. This results in the following model for a measurement $y_i$ of a mouse with age $a$ and treatment $t$:

$$\ln y_i \sim N(\hat{y}_{at}, \sigma_t) \tag{2}$$
$$\hat{y}_{at} = \alpha_a^{age} + \alpha_t^{treatment} + \alpha_{at}^{age:treatment} \tag{3}$$
$$\alpha_a^{age} \sim N(0, 0.3) \tag{4}$$
$$\alpha_t^{treatment} \sim N(0, 0.3) \tag{5}$$
$$\alpha_{at}^{age:treatment} \sim N(0, 0.2) \tag{6}$$
$$\sigma_t \sim HN(0, 0.5) \tag{7}$$

Note that this model does not include any hierarchical parameters because the measured quantities are systemic parameters (e.g., one MAP value per mouse).

#### 2.6 Whisker stimulation responses

As dependent variable we used the ratio of the maximal vessel diameter (after the stimulation) to the pre-stimulation diameter for each mouse at each stage, on a natural logarithmic scale, also known as the "log-change". The latter was standardised by subtracting the overall mean and dividing by the standard deviation, then treated as a single measurement. We used the log-change to facilitate modelling, as log-change is a symmetric and additive measure of relative change [7]. When $v1 - v2 \ll$

1, the log change $\ln \frac{v2}{v1}$ is approximately the same as the more widely used relative difference measure $\frac{v2-v1}{v1}$.

Our final model was a hierarchical multilevel linear model of this quantity with student-T distributed errors (Supplementary Fig. 3). We used Eq. 1 plus the following prior for the student-T degrees of freedom parameter $\nu$, following [6]:

$$\nu \sim Gamma(2, 0.1)$$

We also fit a *big* version of this model which adds interaction effects for vessel-type:treatment and age:treatment and is otherwise identical to the final model. The big model showed no large new effects and slightly worse estimated log predictive density compared with the final model: elpd $349 \pm 40$ vs $353 \pm 40$, elpd difference $3.5 \pm 3.2$.

##### 2.7 Myogenic response

The myogenic response was quantified with the correlation coefficient, CC, between MAP and a vessel's diameter. To simplify modeling, we took inverse hyperbolic tangent function of CC (because CC $\in [-1; 1]$), so that the resulting quantity had support on the entire real number line.

We used Eq. 1, but in this case we also used non-hierarchical prior distributions for treatment effects, as measurements were only available from two treatment types (Supplementary Fig. 4). We also allowed the measurement error parameters $\sigma$ to vary according to vessel type, since this improved model fit and predictive performance.

##### 2.8 RBC velocity and flux

We modelled velocity and flux measurements on natural logarithmic scale as we expected multiplicative effects (Supplementary Fig. 5). Our final model used the normal distribution to describe measurement errors and otherwise followed Eq. 1.

For investigation of interaction effects we fit another *big* model that extended our final model with a vessel-type:treatment interaction effect. For the two models of RBCs velocity, namely *final* and *big*, estimated log-predictive densities (Elpd) were $-207.3 \pm 19.2$ (mean±SEM) and $-207.6 \pm 19.1$ (mean±SEM), respectively, and the estimated elpd difference was $0.36 \pm 1.16$, indicating near-identical performance.

For the two models of RBCs flux, namely *final* and *big*, estimated Elpd were $-181.5 \pm 29.2$ (mean±SEM) and $-182.2 \pm 29.3$ (mean±SEM), respectively,and the estimated elpd difference was $1.42 \pm 0.92$, indicating slightly better predictive performance for the final model than the *big* model. Neither *big* nor *final* model exhibited large interaction effects. Given the very close performance and absence of clear effects, we discarded both *big* models.

##### 2.9 Vessels pulsations $P_d$ and $P_c$

$P_d$ and $P_c$ were estimated by Fourier-transforming time-series of vessels diameters and center positions and estimating the spectral power of the first harmonic at the heartbeat frequency. We didn't analyse higher harmonics as they were typically one order of magnitude lower than the first harmonic. Using only the first harmonic also simplified the modelling.

Because, $P_d$ and $P_c$, being spectral powers, should follow exponential distributions [1], we used exponential generalised linear models for both $P_d$ and $P_c$ (Supplementary Fig. 6). In this model, given measurement $y$ and linear predictor $\eta$ the measurement probability density is given by

$$p(y \mid \eta) = Exponential(y, \lambda) \quad (8)$$

$$= \lambda e^{-\lambda y} \quad (9)$$

$$\ln \frac{1}{\lambda} = \eta \quad (10)$$

The logarithmic link function was chosen so that linear changes in the term $\eta$ induce multiplicative changes in the mean $\frac{1}{\lambda}$ of the measurement distribution, as we believed the effects we wanted to model would be multiplicative. The model otherwise followed Eq. 1.

##### 2.10 Baseline vessel diameter

To model measurements of baseline vessel diameters (Supplementary Fig. 7) we used Eq. 1.

##### 2.11 Topology of collaterals and analysis of number of PSs and bulbs

Measurements of collateral vessel density, tortuosity, length and diameter (Supplementary Fig. 8), and PSs and bulb occurrence rates, were structurally different from the other measurements we analysed, leading to a different statistical modelling approach. Specifically, fewer covariates were available for these data, namely age, mouse id and for branchpoints the branch number, depth and first-order-per-penetrating-arteriole rate. In addition the dependent variables could all be modelled by standard generalised linear model families. We therefore judged that, for these data, it was safe to consider only models that can be specified using the Python library bambi[3]. Whereas Stan allows arbitrary statistical models to be expressed and fitted, bambi allows only models that can be formulated using a formula syntax based on Wilkinson notation[10].

For mouse-level measurements of collateral density and craniotomy diameters we used the following model, represented in bambi's formula language by the two formulae `{y} ~ age` and `sigma ~ age`, where `{y}` is substituted by the name of the log-transformed dependent variable:

$$
\begin{aligned}
y &\sim N(\hat{y}, \sigma) && (11)\\
\hat{y} &= \mu + \alpha^{age} && (12)\\
\mu &\sim N(0, s1) && (13)\\
\alpha^{age} &\sim N(0, s2) && (14)\\
\ln \sigma &\sim N(0, s3) && (15)
\end{aligned}
$$

For collateral-vessel-level measurements of tortuosity, length and diameter, we used a model represented in bambi's formula syntax as `{y} ~ (1|mouse_id) + (1|age)`. Again `y` is substituted by the name of the log-transformed dependent variable. For tortuosity measurements, we subtracted the theoretical minimum value 1 before log-transforming the measurements. The full model specification is as follows:

$$
\begin{aligned}
y &\sim N(\hat{y}, \sigma) && (16)\\
\hat{y} &= \mu + \alpha^{age} + \alpha^{mouse} && (17)\\
\mu &\sim N(0, s1) && (18)\\
\alpha^{age} &\sim N(0, t1) && (19)\\
\alpha^{mouse} &\sim N(0, t2) && (20)
\end{aligned}
$$

For modeling PSs and bulb occurrence rates we used a logistic regression model with bambi formula `{y}['True'] ~ age + branch_number + ln_depth + logit_firstorder_per_pa`. In this case the dependent variable is a boolean indicating whether the given vessel branching point had a sphincter or bulb. The full model is as follows:

$$y \sim bernoulli(logit^{-1}\eta) \tag{21}$$

$$\eta = \mu + \alpha^{age} + \beta^{branch} \cdot branch + \beta^{depth} \cdot \ln depth + \beta^{first\ order} \cdot logit(first\ order\ rate) \tag{22}$$

$$\mu \sim N(0, s1) \tag{23}$$

$$\beta^{branch} \sim N(0, s2) \tag{24}$$

$$\beta^{depth} \sim N(0, s3) \tag{25}$$

$$\beta^{first\ order} \sim N(0, s4) \tag{26}$$

The prior hyperparameters $s_i, t_i$ were the bambi defaults, which aim to be weakly informative while automatically scaling to match the predictors.

#### 2.12 Capillary topology

Here we describe modeling of capillary density, length, and tortuosity. We modeled capillary density measurements on natural logarithmic scale as we expected multiplicative effects. In this case we noticed that adjacent vessel types tended to have similar average measurement values, making the independent model in our common structure inappropriate. We therefore constructed a model that smooths the prior distributions of measurements from adjacent vessel types using a random walk prior (Supplementary Fig. 9). Since we were interested in whether the overall pattern of average measurements by vessel-type would differ for old and adult mice, we modelled age and vessel type jointly using the following equations:

$$\alpha_{a,v}^{\text{age, vessel type}} \sim \begin{cases} N(0,1), v = 1 \\ N(\alpha_{a,v-1}^{\text{age, vessel type}}, \lambda_a^\alpha), v > 1 \end{cases} \tag{27}$$

In this equation, $\lambda^\alpha$ is a non-negative parameter representing the degree of smoothness of the random walk in each age category. We also used a Gaussian random walk prior for the model's measurement error parameter $\sigma$:

$$\ln \sigma_v \sim \begin{cases} N(0, sd(\ln y), v = 1 \\ N(\ln \sigma_{v-1}, \lambda^s), v > 1 \end{cases} \tag{28}$$

This approach to smoothing parameters corresponding to ordered categories is essentially the same as that Gao et al. [4] used to model age effects on voting behaviour. As explained in that paper, the random walk priors allow for information sharing between categories, without the need for detailed assumptions about the functional form of the overall relationship.

The model was otherwise the same as Eq. 1.

Similarly to how we analysed capillary density, we used random-walk priors to smooth adjacent vessel type effects within age categories and measurement errors for adjacent vessel types (Supplementary Fig. 9). In order to appropriately capture the increased heterogeneity in steps between average measurements for this data type, we modelled the random walk with student-t distributions, with the degrees of freedom parameters modelled as in the whisker stimulation analysis.

To analyse capillary length, we again used random-walk priors to smooth adjacent vessel type effects within age categories and measurement errors for adjacent vessel types, with the random walk modelled by student-t distributions (Supplementary Fig. 9).
